# Artemisinin resistance mutations in *Pfcoronin* impede hemoglobin uptake

**DOI:** 10.1101/2023.12.22.572193

**Authors:** Imran Ullah, Madeline A. Farringer, Anna Y. Burkhard, Erica Hathaway, Malhar Khushu, Bailey C. Willett, Sara H. Shin, Aabha I. Sharma, Morgan C. Martin, Kairon L. Shao, Jeffrey D. Dvorin, Daniel L. Hartl, Sarah K. Volkman, Selina Bopp, Sabrina Absalon, Dyann F. Wirth

**Author notes:** Corresponding authors: Dyann F. Wirth, Sabrina Absalon.

## Abstract

Artemisinin (ART) combination therapies have been critical in reducing malaria morbidity and mortality, but these important drugs are threatened by growing resistance associated with mutations in *Pfcoronin* and *Pfkelch13*. Here, we describe the mechanism of *Pfcoronin*-mediated ART resistance. *Pf*Coronin interacts with *Pf*Actin and localizes to the parasite plasma membrane (PPM), the digestive vacuole (DV) membrane, and membrane of a newly identified preDV compartment—all structures involved in the trafficking of hemoglobin from the RBC for degradation in the DV. *Pfcoronin* mutations alter *Pf*Actin homeostasis and impair the development and morphology of the preDV. Ultimately, these changes are associated with decreased uptake of red blood cell cytosolic contents by ring-stage *Plasmodium falciparum*. Previous work has identified decreased hemoglobin uptake as the mechanism of *Pfkelch*13-mediated ART resistance. This work demonstrates that *Pf*Coronin appears to act via a parallel pathway. For both *Pfkelch13*-mediated and *Pfcoronin*-mediated ART resistance, we hypothesize that the decreased hemoglobin uptake in ring stage parasites results in less heme-based activation of the artemisinin endoperoxide ring and reduced cytocidal activity. This study deepens our understanding of ART resistance, as well as hemoglobin uptake and development of the DV in early-stage parasites.

## Introduction

Rapidly acting and potent antimalarial drugs, such as artemisinin (ART) and others in its class, are critical components of current front-line treatments, and these drugs have contributed to yearly reductions in malaria mortality for the past two decades^1^. However, partial resistance to ART and artemisinin combination therapies (ACTs) threatens their efficacy^2,3^. ART treatment failures initially emerged in Southeast Asia, and were associated with mutations in the *Plasmodium falciparum kelch13* locus^4^. The mechanism of *Pfkelch13*-mediated ART resistance is well studied, and prior work has demonstrated reduced hemoglobin uptake in ART-resistant *Pfkelch13* mutant parasites^5,6^. ART is a sesquiterpene lactone with an endoperoxide bridge that must be cleaved to potentiate antimalarial activity. Typically, this activation occurs via heme^7^, likely derived via endocytosis and digestion of hemoglobin. Thus, the observed reduction in endocytosis in *Pfkelch13* mutants may decrease the availability of free heme, slowing the activation of ART and leading to ART resistance ^5,6^.

As part of our previous efforts to understand potential ART-resistance mechanism(s) in Africa, we demonstrated *Pfcoronin* to be a major driver of *in vitro* partial ART resistance in Senegalese isolates of *P. falciparum*^8,9^. Coronin proteins are widely conserved, and while many organisms have multiple *coronin* genes, *P. falciparum* and other unicellular pathogens encode just one^10,11^. Like other Coronin orthologs, *Pf*Coronin includes a WD40 domain. These domains are thought to facilitate protein-protein interactions and are involved in a wide range of cellular functions^12,13^. Coronin proteins have been associated with actin-binding activities in multiple organisms^14^; in the apicomplexan parasite, *Toxoplasma gondii, Tg*Coronin regulates the rate and extent of actin polymerization, and is required for endocytosis and membrane recycling^15^. In *P. berghei, Pb*Coronin is implicated in the regulation of gliding motility in sporozoites^16^. In *P. falciparum, in vitro* biochemical studies have confirmed that the WD40 domain of *Pf*Coronin binds actin and organizes F-actin filaments into parallel bundles^17^. In late asexual blood-stage parasites, *Pf*Coronin localizes to the parasite periphery, potentially linking actin filaments to the plasma membrane^17^; localization in earlier stages has not been previously assessed.

Here, we study *Pf*Coronin and identify its interaction partners in both rings and late-stage parasites. Furthermore, we assess the impact of *Pfcoronin* mutations on protein levels, subcellular localization, and hemoglobin uptake to improve our understanding of potential early-stage parasite DV development and ART activation.

## Results

### Reduced interaction of *Pf*Coronin mutants with *Pf*Actin in rings

Previously, we demonstrated that mutations in the *Pfcoronin* gene are a major driver of *in vitro* evolved ART resistance in Senegalese isolates of *P. falciparum*, as measured by the ring-stage survival assay (RSA)^8^. Either a single G50E mutation (Senegalese isolate from Thiès) or paired R100K/E107V mutations (Senegalese isolate from Pikine) are sufficient to confer resistance^8,9^. *Pf*Coronin contains a seven-bladed WD40 domain with Gly50 located in the first blade, and both Arg100 and Glu107 located in the second blade. We focused on the Pikine *P. falciparum* isolate in this study. WD40 domains have previously been associated with the facilitation of protein-protein interactions^12,13^. Therefore, to identify interaction partners of *Pf*Coronin, we appended the spaghetti monster-human influenza hemagglutinin (HA) tag (smHA)^18^ to the c-terminus of the endogenous *Pfcoronin* locus in both *Pfcoronin*^*WT*^ and *Pfcoronin*^*R100K/E107V*^ genetic backgrounds (Supplementary Fig. 1). We immunoprecipitated (IP) *Pf*Coronin-smHA protein in both early ring and later trophozoite-stage parasites [3-9 or 32 – 40 hours post-invasion (hpi)] using anti-HA antibodies (Supplementary Fig. 2a); we also included an untagged *Pfcoronin*^*WT*^ control line. Co-purified proteins were identified by liquid chromatography with tandem mass spectrometry (LC-MS/MS). To identify potential *Pf*Coronin interacting partners, we first filtered out proteins that were found in the untagged line and normalized prey protein abundance to that of *Pf*Coronin bait abundance [(Supplementary Data 1 (3-9 hpi); Supplementary Data 2 (32 – 40 hpi)]. We established two criteria to prioritize top *Pf*Coronin interacting partners: (1) the protein must be detected in all replicates of either *Pf*Coronin^WT^-smHA or *Pf*Coronin^R100K/E107V^-smHA parasites and (2) its abundance must be at least 5% of *Pf*Coronin (bait) protein abundance. To investigate interactions that were potentially affected by mutations in *Pfcoronin*, we looked for differentially-associated proteins with, greater than 5x change in normalized abundance between *Pf*Coronin^WT^-smHA and *Pf*Coronin^R100K/E107V^-smHA, and/or for which the *p*-value (unpaired t-test between *Pf*Coronin^WT^-smHA and *Pf*Coronin^R100K/E107V^-smHA) was less than 0.05 [(Supplementary Fig. 2b and Supplementary Data 3 (3-9 hpi); Supplementary Fig. 2c and Supplementary Data 4 (32 – 40 hpi)]. Interestingly, in both stages, the major interacting partner of *Pf*Coronin was *Pf*Actin (Fig. 1a-b), consistent with its role in regulating actin dynamics. At ring stage, this interaction was reduced by three-fold in *Pf*Coronin^R100K/E107V^-smHA parasites (*p* = 0.0113; Fig. 1a and Supplementary Fig. 2b; Supplementary Data 3); this reduction did not occur in the late-stage parasites (*p* = 0.4787; Fig. 1b and Supplementary Fig. 2c; Supplementary Data 4). Additionally, while polyubiquitin (*Pf*Ub) was an abundant interaction partner of *Pf*Coronin^WT^-smHA in ring-stage parasites (3-9 hpi), its interactions with *Pf*Coronin^R100K/E107V^ were dramatically reduced by 15-fold (*p*=0.0282; Fig. 1a and Supplementary Fig. 2b; Supplementary Data 3). In late-stage parasites, we did not observe substantial or significant changes in associations with *Pf*Ub by *Pf*Coronin^WT^-smHA vs. *Pf*Coronin^R100K/E107V^-smHA (*p* = 0.4493; Supplementary Fig. 2c and Supplementary Table 4).

**Fig. 1:**
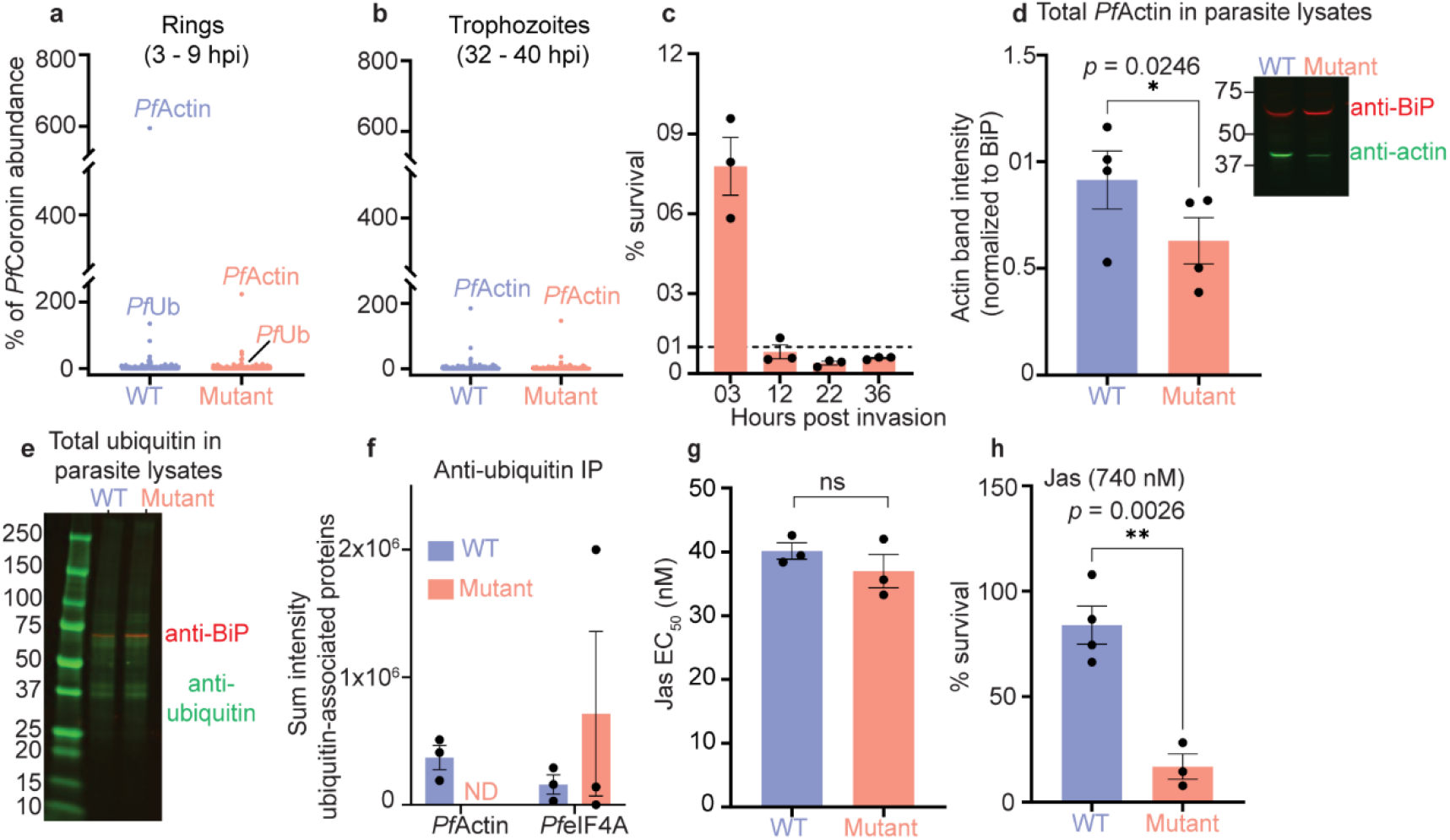
Mutations in *Pfcoronin* alter *Pf*Actin homeostasis. (a) Immunoprecipitation of *Pf*Coronin-smHA and associated proteins in ring and (b) late trophozoite-stage parasites (n=4 rings; n = 6 trophozoites). (c) RSA survival (%) following 700 nM DHA pulse for 6-hour increments at various points in the intraerythrocytic developmental cycle. (d) Total *Pf*Actin levels in lysates from parasites expressing *Pf*Coronin^R100K/E107V^-smHA (mutant) were compared by western blot to lysates from parasites expressing *Pf*Coronin^WT^-smHA (representative blot shown). Blots were probed with anti-actin (mouse monoclonal; green) and anti-BiP (loading control; rabbit polyclonal, red) antibodies. Bar graph shows densitometric quantification of *Pf*Actin, normalized against *Pf*BiP. (e) Global ubiquitination in lysates from parasites expressing *Pf*Coronin^WT^-smHA and *Pf*Coronin^R100K/E107V^-smHA (mutant) were compared by western blot (representative blot shown). Blots were probed with an anti-ubiquitin (mouse mAb FK2, green) and anti-BiP (loading control; rabbit polyclonal red) antibodies. (f) Immunoprecipitation of anti-*Pf*Ub associated proteins. Complete results for all proteins are shown in Supplementary Data 5. Highly consistent proteins, i.e., those identified in all replicates of either *Pf*Coronin^R100K/E107V^-smHA or *Pf*Coronin^WT^-smHA, are shown in Supplementary Fig. 4c. (g) E_50_ values of JAS and (h) percent survival of parasites treated for 6 hrs., starting at 3 hpi, with 740 nM JAS.

### *Pfcoronin* mutations only confer ART resistance in rings

Overall, comparison between *Pf*Coronin interaction partners in *Pf*Coronin^WT^ vs. *Pf*Coronin^R100K/E107V^ parasites showed far more differences at ring-stage than in late-stage parasites. Interestingly, this corresponds to the timing of ART clinical resistance, which manifests as delayed clearance of early ring-stage parasites. Additionally, *Pfkelch13*-mediated ART resistance in cultured parasites is only evident at the ring stage^4,6^. As a result, the ring-stage survival assay (RSA), where ring-stage parasites are exposed to physiological concentrations of dihydroartemisinin (DHA) for just 6 hours, has become the standard assay for *in vitro* ART resistance, with greater than 1% survival indicating ART resistance^4^. To assess whether *Pfcoronin*-mediated ART resistance is stage-specific, we treated parasites with pulses of 700 nM DHA for 6-hour increments across development. Similarly, to *Pfkelch13* mutants, *Pfcoronin*-mediated artemisinin resistance is ring-stage specific: *Pfcoronin*^*R100K/E107V*^ mutants did not show any resistance phenotype beyond the first 6 hours of the lifecycle (Fig. 1c).

### *Pfcoronin* mutations reduce *Pf*Actin protein levels in rings

Since we observed reduced interactions with *Pf*Actin in *Pfcoronin*^*R100K/E107V*^ mutant ring-stage parasites, we were interested in whether these changes represented global differences in *Pf*Actin protein levels. We performed western blot analysis on ring-stage parasite lysates to assess whether *Pf*Actin abundance differs between *Pfcoronin*^*WT*^ and *Pfcoronin*^*R100K/E107V*^ mutant parasites. We found that *Pfcoronin*^*R100K/E107V*^ parasites contained 31 ± 5.8 % (mean ± SEM) less *Pf*Actin, compared to *Pfcoronin*^*WT*^ parasites in rings (*p* = 0.027; Fig. 1d; see Supplementary Fig. 3a for full western blot images and additional examples). This decrease in *Pf*Actin protein levels is consistent with the reduced interactions with *Pf*Actin in *Pfcoronin*^*R100K/E107V*^ mutant ring-stage parasites; however, it is not clear how *Pfcoronin*^*R100K/E107V*^ mutations mediate these changes. We did not observe changes in *Pf*Actin protein levels in late-stage parasites (Supplementary Fig. 3b).

### *Pf*Actin is ubiquitinated only in *Pfcoronin*^*WT*^ parasites in rings

To gain further insights into the changes associated with *Pfcoronin*^*R100K/E107V*^ mutations, we focused on the second most abundant *Pf*Coronin interaction partner, annotated as polyubiquitin, which exhibited striking 15-fold changes in associations with *Pf*Coronin^R100K/E107V^-smHA, compared to *Pf*Coronin^R100K/E107V-smHA^, in ring stage parasites (Fig. 1a, Supplementary Fig. 2b; Supplementary Data 3). We questioned whether the dramatic reduction in apparent interactions with “polyubiquitin” in *Pfcoronin*^*R100K/E107V*^ mutant ring-stage parasites represented global differences in ubiquitination or specific changes in ubiquitination of *Pf*Coronin or its interaction partners.

First, to explore overall protein ubiquitination in *Pfcoronin*^*WT*^*-smHA* vs. *Pfcoronin*^*R100K/E107V*^*-smHA* parasites, we generated ring-stage parasite lysates (3-9 hpi) and performed anti-ubiquitin Western blots. We found no noticeable difference between the *Pfcoronin*^*WT*^*-smHA* and the *Pfcoronin*^*R100K/E107V*^*-smHA* mutant (Fig. 1E; see Supplementary Fig. 4a for full western blot images and additional examples), suggesting that global changes in the ubiquitination system were unlikely. To identify potential changes in specific ubiquitinated proteins, we identified ubiquitin-interacting partners using an anti-ubiquitin antibody in *Pfcoronin*^*WT*^*-smHA* and *Pfcoronin*^*R100K/E107V*^*-smHA* ring-stage parasites (Supplementary Fig. 4b). As before, we focused on proteins that were detected in all replicates of either *Pfcoronin*^*WT*^*-smHA* or *Pfcoronin*^*R100K/E107V*^*-smHA* parasites (Supplementary Fig. 4c and Supplementary Data 5). Of these putatively-ubiquitinated proteins, only two were found to interact with *Pf*Coronin in this study (Supplementary Data 3): eukaryotic initiation factor 4A (eIF4A) and *Pf*Actin. *Pf*Actin was of particular interest because of its high abundance in anti-smHA (*Pf*Coronin) IP experiments and its known interaction with *Pf*Coronin. Interestingly, associations between *Pf*Actin and ubiquitin were detected in *Pfcoronin*^*WT*^*-smHA* parasites, but not in *Pfcoronin*^*R100K/E107V*^*-smHA* samples (Fig. 1f).

Detection of *Pf*Actin via anti-ubiquitin antibodies could indicate that *Pf*Actin itself is ubiquitinated, or that *Pf*Actin physically associates with another protein (or proteins) that is ubiquitinated. To distinguish between these possibilities, we performed Lys-ϵ-Gly-Gly (diGly) proteomics, which identifies not only which proteins are ubiquitinated, but also at which sites^19^ (Supplementary Fig. 5a). We identified ubiquitination events at K114 and K62 of *Pf*Actin only in *Pfcoronin*^*WT*^ parasites (Supplementary Fig. 5b; Supplementary Data 6).

### *Pfcoronin* mutants sensitize *Pf*Actin to jasplakinolide

Our data revealed that *Pfcoronin*^*R100K/E107V*^*-smHA* parasites have changes in *Pf*Actin abundance and ubiquitination. We therefore hypothesized that *Pfcoronin* mutations lead to defects or dysregulation of *Pf*Actin dynamics. We reasoned that these changes might alter sensitivity to jasplakinolide (JAS), which stabilizes *Pf*Actin filaments and promotes polymerization, ultimately disrupting normal *Pf*Actin dynamics. We tested this hypothesis using *Pfcoronin*^*WT*^ and *Pfcoronin*^*R100K/E107V*^ parasites (DHA selected Pikine_R line)^8^. We first estimated sensitivity to JAS using a standard 72 h growth assay and found no difference between *Pfcoronin*^*WT*^ (EC_50_ = 40.1 ± 1.3 nM) and *Pfcoronin*^*R100K/E107V*^ parasites (EC_50_ = 37 ± 2.6 nM; Fig. 1g and Supplementary Fig. 5c). To assess possible differences in ring-stage parasites, we performed RSA-type assays on *Pfcoronin*^*WT*^ and *Pfcoronin*^*R100K/E107V*^ parasites. Parasites were treated for 6 hrs, starting at 3 hpi, with 740 nM JAS (∼20X EC_50_). JAS had a modest effect on *Pfcoronin*^*WT*^ parasite survival (84 ± 9% survival) but significantly reduced the survival of *Pfcoronin*^*R100K/E107V*^ parasites (17 ± 6 % survival) (Fig. 1h). Thus, *Pfcoronin* mutations sensitize parasites to *Pf*Actin disruptions, suggesting some level of underlying *Pf*Actin dysfunction or dysregulation, particularly in early ring-stage parasites.

### *Pf*Coronin localizes to the periphery during late schizogony

We used immunofluorescence assay (IFA) to investigate *Pf*Coronin localization. Previous IFA studies^17^ have shown that *Pf*Coronin is localized to the membrane of individual merozoites during late schizogony. Consistent with these studies, *Pf*Coronin-smHA was localized to the merozoite membrane. Potential co-localization with the major interacting partner, *Pf*Actin, was also assessed. *Pf*Actin was distributed throughout the parasite (Fig. 2) and was found in close proximity to *Pf*Coronin at the membrane, but its localization did not substantially overlap with *Pf*Coronin (Fig. 2). We did not find any observable differences in localization of *Pf*Coronin and *Pf*Actin in WT vs. mutant late stage, segmenting parasites (Fig. 2).

**Fig. 2:**
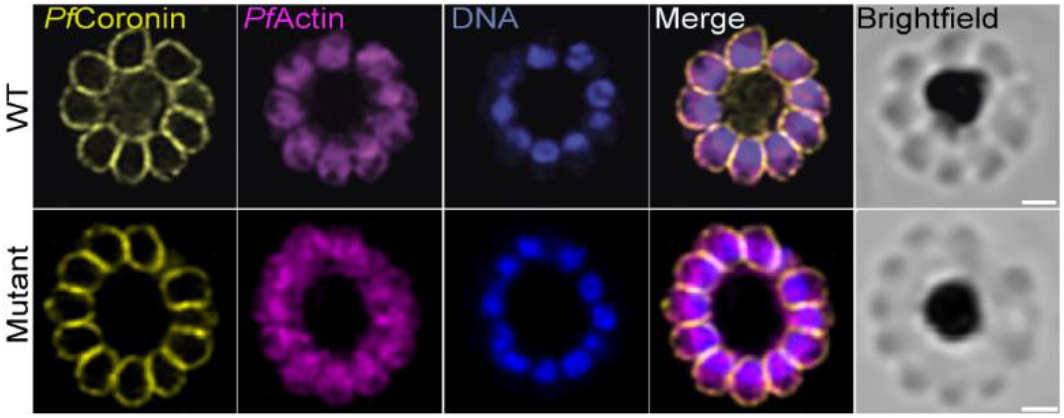
Immunofluorescent characterization of *Pf*Coronin and *Pf*Actin localization. Super-resolution fluorescence microscopy using Zeiss LSM980 with Airyscan2 shows localization of *Pf*Coronin and *Pf*Actin in segmented schizonts. Scale bars = 2 μm.

### *Pf*Coronin marks vesicular ultrastructures across development

To gain further insights into *Pf*Coronin localization, we performed ultrastructure expansion microscopy (U-ExM) imaging using Airyscan2 super-resolution microscopy^20–22^. *Pf*Coronin had not previously been examined during ring-stage—the most relevant stage for ART resistance. We therefore assessed *Pf*Coronin^WT^ localization in rings as well as throughout development (Fig. 3). *Pf*Coronin was found at the membranes, likely representing the parasite plasma membrane (PPM) and/or parasite vacuolar membrane (PVM) (Fig. 3). In rings (6 – 15 hpi), in addition to PPM/PVM, *Pf*Coronin is localized around an intracellular vacuole that may be a precursor to the DV; we refer to this vacuole as the <u>“pre-DV compartment” (PreDV)</u>. The identification of preDV-like vacuoles was first observed in earlier research utilizing electron microscopy (EM)^23^. PreDV contains host cytoplasmic content, predominantly hemoglobin, demonstrated by staining with NHS ester, (Fig. 3, rings) and further confirmed through staining of biotin-dextran (RBCs had been pre-loaded with biotin-dextran; see below; Supplementary Fig. 6). Further investigation in trophozoite stage parasites (26 – 34 hpi) confirmed localization of *Pf*Coronin at the DV membrane (Fig. 3). Interestingly, in addition to the PPM/PVM and DV membrane, *Pf*Coronin is localized to small double and single membrane-bound vacuoles (Fig. 3). These small vacuoles are located near the cytostome but do not overlap with it (Fig. 3).

**Fig. 3:**
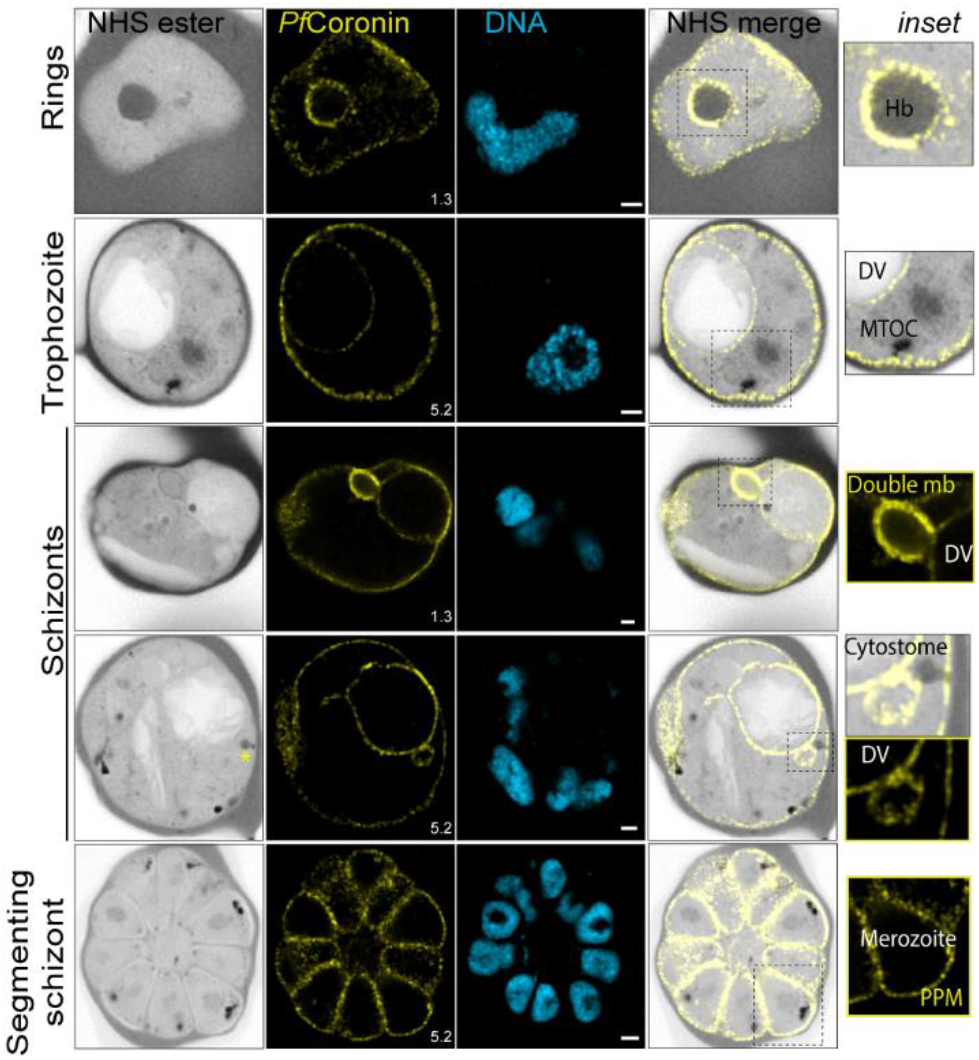
*Pf*Coronin marks ultrastructures in WT parasites throughout blood stage development. U-ExM shows *Pf*Coronin localization throughout the intraerythrocytic cycle in WT parasites. Cytostome is marked with a yellow asterisk in the (NHS ester; schizonts). Alexa Fluor 405-conjugated NHS ester (protein density), HA (*Pf*Coronin, yellow), Sytox deep red (DNA, Cyan). Images are maximum-intensity projections; number on image = Z-axis thickness of projection in μm. Scale bars = 2 μm.

Cytostomes, double-membrane invaginations visible at the parasite periphery, are characterized by relatively small structures with a narrow neck connected to a wider body protruding into the parasite cytosol. Notably, *Pf*Kelch13 localizes to the neck of the cytostome. In our later-stage U-ExM localization, direct association between *Pf*Coronin and the cytostomal body itself is not observed (Fig. 3). However, considering its presence at PPM/PVM, its involvement at the collar of the cytostome cannot be entirely ruled out. In addition, we observed *Pf*Coronin-coated interconnected chains of small and large vacuoles (Supplementary Fig. 7). IFA analysis of early schizonts showed similar localization to the PPM/PVM, DV membrane, and smaller vesicles near the DV (Supplementary Fig. 8).

### *Pf*Coronin mutations disrupt its ultrastructure localization and protein levels

We next compared localization of *Pf*Coronin^WT^-smHA vs. *Pf*Coronin^R100K/E107V^-smHA using U-ExM. Interestingly, *Pf*Coronin^R100K/E107V^ formed disorganized puncta at ring stage, and was not localized at the PPM/PVM, DV, or preDV, but shows a WT-like normal distribution in late schizonts (Fig. 4a; Supplementary Fig. 9). We hypothesized that the disorganized punctate localization of *Pf*Coronin^R100K/E107V^-smHA might be associated with reduced protein levels and therefore we performed immunoblot-based quantification analysis. We found significantly less *Pf*Coronin protein in the R100K/E107V mutant than in WT ring-stage parasites (3-9 hpi), but protein levels were unchanged in late-stage parasites (32-40 hpi) (Fig. 4b; see Supplementary Fig. 10 for full western blot images and additional examples). This result suggests that *Pfcoronin* mutations result in reduced protein levels.

**Fig. 4:**
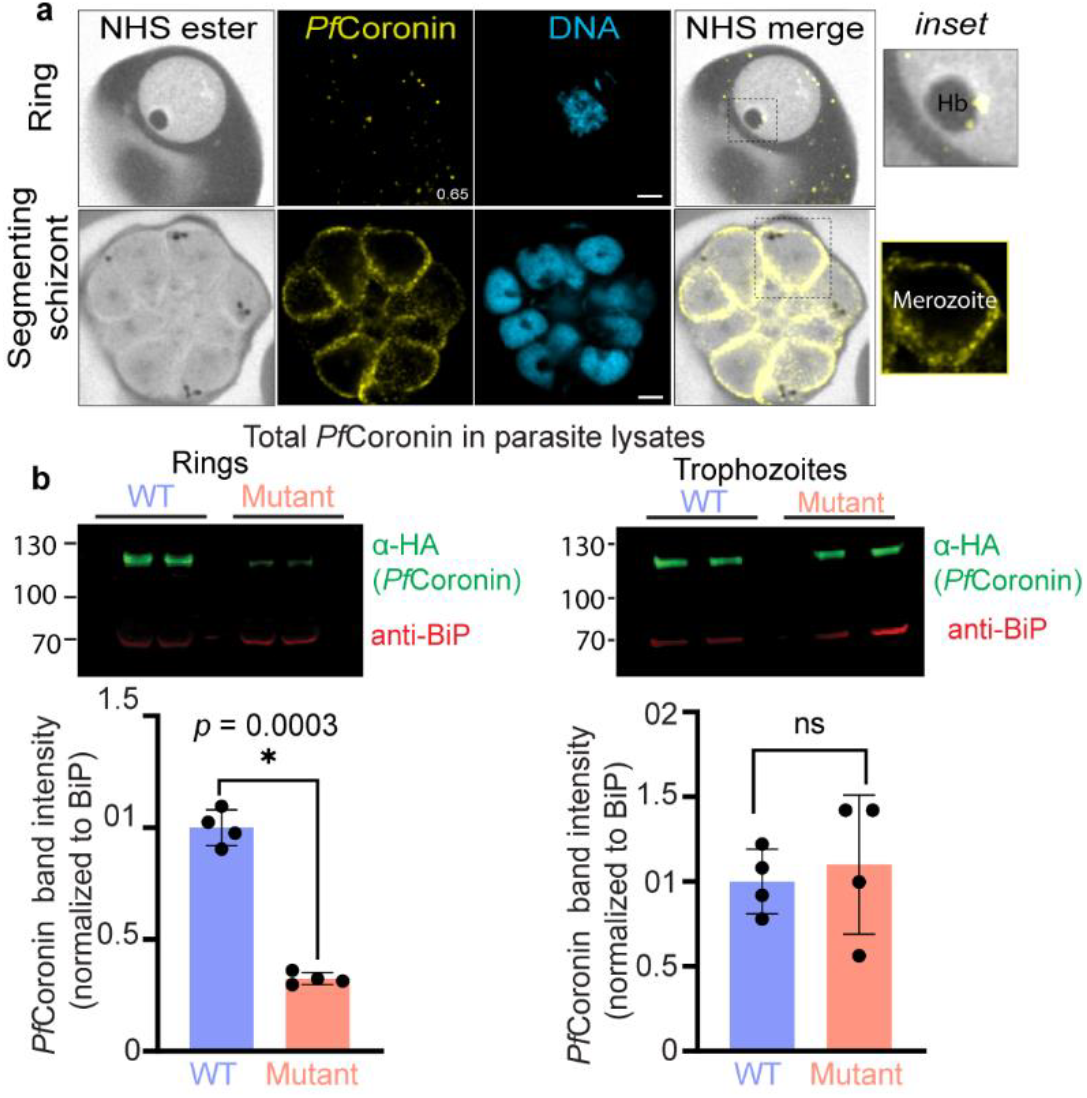
*Pfcoronin* mutations impact localization and protein levels in rings. (a) U-ExM shows *Pf*Coronin^R100K/E107V^-smHA (mutant) localization in rings and late schizont stage parasites. Alexa Fluor 405-conjugated NHS ester (protein density), HA (*Pf*Coronin, yellow), Sytox deep red (DNA, Cyan). Images are maximum-intensity projections, number on image = Z-axis thickness of projection in μm. Scale bars = 2 μm. (b) Total *Pf*Coronin levels in lysates from parasites expressing *Pf*Coronin^R100K/E107V^-smHA (mutant) were compared to lysates from parasites expressing *Pf*Coronin^WT^-smHA via western blot (representative blot shown). Blots were probed with anti-HA (*Pf*Coronin; mouse monoclonal; green) and anti-BiP (loading control; rabbit polyclonal, red) antibodies. Bar graph shows densitometric quantification of *Pf*Coronin, normalized against *Pf*BiP.

### Mutations affect the size and number of *Pf*Coronin-coated hemoglobin-filled preDV and invaginations in early rings

The *Pf*Coronin-coated preDV, which houses host cell cytoplasmic contents, e.g., hemoglobin (Fig. 3, rings; Supplementary Fig. 6), is visible in many ring-stage parasites. *Pf*Coronin is also localized around large invaginations in early rings (Fig 5a). While the preDV appears as a fully-contained vacuole in older ring-stage parasites (>9hpi), invaginations are more commonly observed in early rings (<9hpi; Fig 5a). In our U-ExM studies of early ring-stage parasites (0-9 hpi), the *PfCoronin*^*R100K/E107V*^*-smHA* mutant parasites showed a significant decrease in the presence of the preDV and/or invagination, with only 23.8% of parasites displaying this feature (Fisher’s exact test, *p* < 0.0001; Supplementary Fig. 11a). In contrast, all analyzed images of *Pfcoronin*^*WT*^*-smHA* parasites showed the presence of the preDV and/or invagination. We measured the size of preDV /invagination and normalized it to the parasite size for both *Pfcoronin*^*WT*^*-smHA* and *Pfcoronin*^*R100K/E107V*^*-smHA* mutant parasites. We observed a 21% reduction in the preDV /invagination in *Pfcoronin*^*R100K/E107V*^*-smHA* parasites (unpaired t-test; *p* = 0.0340; Fig 5b; Supplementary Fig.11b-d). These observations led us to hypothesize that *Pf*Coronin mutations might impair hemoglobin uptake, possibly via abnormal development of the preDV.

**Fig. 5:**
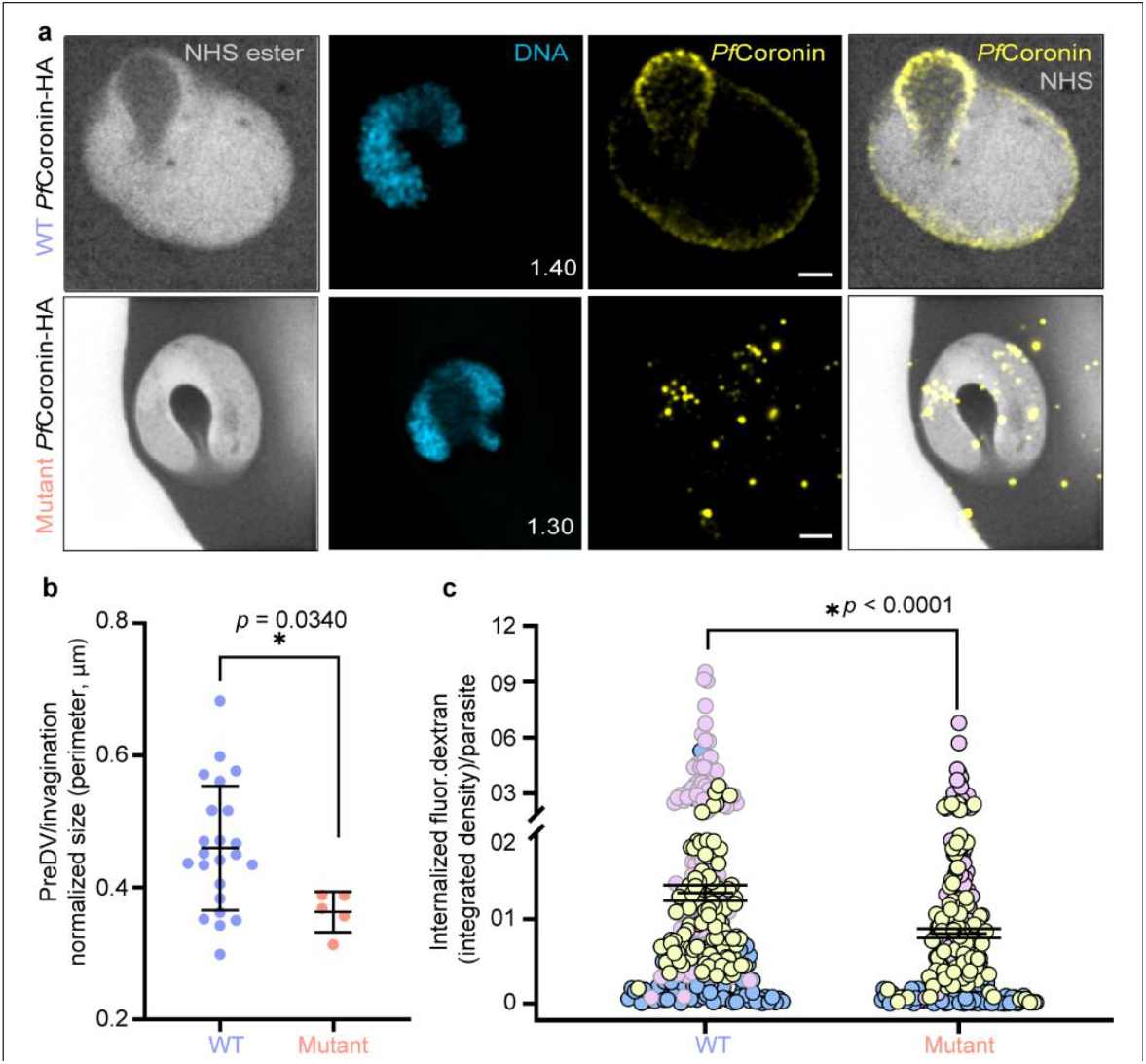
*Pfcoronin* mutations reduce invagination size and host cell content uptake. (a) Representative U-ExM images of *Pf*Coronin-labeled invaginations in *Pfcoronin*^*WT*^*-smHA* and *Pfcoronin*^*R100K/E107V*^*-smHA* mutant parasites. Alexa Fluor 405-conjugated NHS ester (protein density), HA (*Pf*Coronin, yellow), Sytox deep red (DNA, cyan). Images are maximum-intensity projections, with the number on the image indicating the Z-axis thickness of the projection in μm. Scale bars = 2 μm. (b) Comparison of invagination perimeter in WT (n=23) vs. mutant (n=5) parasites, normalized against parasite perimeter. See Supplementary Fig. 11d for parasite size measurements. *p*-value (unpaired t-test) and % reduction of the mean are indicated. (c) Comparison of internalized fluorescein-dextran in WT vs. mutant (R100K/E107V) *Pfcoronin* parasites by integrated density (vesicle size + intensity). WT *Pfcoronin*; n = 286, mutant *Pfcoronin*; n = 286. Each data point represents fluorescein-dextran uptake by a single parasite. *p*-values (unpaired, nonparametric Mann-Whitney test) is indicated. These data were pooled from the individual biological replicates shown in Supplementary Fig. 14.

### Mutations in *Pfcoronin* reduce hemoglobin uptake

Prior studies demonstrated that the uptake of RBC contents is decreased in ART-resistant *Pfkelch13* mutants^5,6^. This reduction in endocytosis may slow the activation of ART by decreasing the availability of free heme. Localization of *Pf*Coronin to the membranes of the parasite periphery, vesicular structures, including internalized preDV and invaginations, and DV—all structures involved in the trafficking of hemoglobin from the RBC to the DV—suggested that *Pf*Coronin, similarly to *Pf*Kelch13, may play a role in the uptake of hemoglobin and other RBC contents. While mutations in *Pfcoronin* reduced the number and size of the preDV, it led us to hypothesize that *Pfcoronin* mutations might impair hemoglobin uptake, possibly via abnormal development of the preDV. To test this hypothesis, we performed a previously published microscopy-based endocytosis assay, which uses fluorescein-dextran as a tracer for uptake of host cell contents^5^. Fluorescein-dextran-loaded RBCs were mixed with late schizonts and incubated for 3 hours to allow invasion. After 3 hours, the remaining schizonts were removed by sorbitol synchronization. The resulting population of 0-3 hpi parasites were allowed to grow within the fluorescein-dextran-loaded RBCs for 6 hours. Following this incubation, RBC cell contents were removed with saponin, leaving just the young parasites, which were fixed. Image analyses were performed as previously described^5^ (Supplementary Fig. 12a-b). Since *Pfkelch13*^*C580Y*^ mutants had previously been assessed in the same assay^6^, we used *Pfkelch13*^*C580Y*^ mutants^9^ to validate the assay. We measured a significant 34.6% reduction in fluorescein-dextran uptake in *Pfkelch13*^*C580Y*^ mutant parasites compared to the *Pikine*^*WT*^ parent (unpaired Mann-Whitney test; *p* < 0.0001; Supplementary Fig. 13a; see supplementary Fig. 13b-d for individual bioreps). This figure closely matches the 37.47% reduction previously reported in *Pfkelch13*^*C580Y*^ mutants^5^. We then assessed fluorescein-dextran uptake in *Pfcoronin*^*R100K/E107V*^ mutants. Interestingly, we observed a 36.1% reduction in fluorescein-dextran uptake in *Pfcoronin*^*R100K/E107V*^ mutants compared to WT parasites (unpaired Mann-Whitney test; *p* < 0.0001; Fig. 5c; see Supplementary Fig. 14). In summary, our findings reveal that both *Pfcoronin* and *Pfkelch13* alter endocytosis and reduce hemoglobin uptake to mediate ART resistance. However, the unique localization of *Pf*Coronin to the parasite PVM/PPM, preDV, invagination, and the DV membrane, along with its impact on the number and size of invaginations, suggests that *Pf*Coronin facilitates hemoglobin uptake through a different mechanism.

## Discussion

In this work, we provide insights into the molecular mechanism of *Pfcoronin*-mediated ART resistance. We demonstrate that *Pfcoronin* mutations confer ART resistance only in early ring-stage parasites. In line with the ring-stage specific phenotype, associations of *Pf*Coronin with the two major interacting partners, *Pf*Actin and polyubiquitin, are significantly reduced in *Pfcoronin* mutants at ring stage, but not at later stages. Additionally, *Pf*Actin protein levels are reduced in ring-stage *Pfcoronin*^*R100K/E107V*^ mutants, compared to *Pfcoronin*^*WT*^ parasites, and ring-stage *Pfcoronin*^*R100K/E107V*^ mutants are sensitized to killing by JAS, which promotes actin polymerization. U-ExM revealed that *Pf*Coronin is localized to the PVM/PPM and the DV membrane, as well as to a previously undescribed hemoglobin-filled preDV, visible as an invagination connected to the RBC cytosol in early rings, and entirely contained within the parasite in older ring-stage parasites. Mutations in *Pfcoronin* alter the number and size of the preDVs. Finally, we found that *Pfcoronin* mutations significantly reduce uptake of RBC contents. Similar reductions in hemoglobin uptake have been associated with *Pfkelch13*-mediated ART resistance^5^.

We present multiple lines of evidence indicating that mutations in *Pfcoronin* impact *Pf*Coronin protein levels, its subcellular localization, and homeostasis of *Pf*Actin—the major interacting partner of *Pf*Coronin—in ring-stage parasites. Actin filament polymerization and decay enable the formation of specialized structures and drive multiple complex cellular processes such as vesicular trafficking and endocytosis^24^. In *P. falciparum*, dysregulation of *Pf*Actin using the filament stabilizing agent, JAS, alters cytostome morphology and inhibits hemoglobin trafficking to the DV^25^. We find that mutations in *Pfcoronin* alter the regulation of *Pf*Actin, including changes in quantity and in ubiquitination status. These findings are consistent with a previous study in *P. chabaudi*, where actin was among one of three proteins identified to be ubiquitinated in ring-stage parasites^26^. Studies in other organisms have also reported ubiquitination of actin^27^, and ubiquitin has been shown to play a role in actin homeostasis^28^. In *Pfcoronin*^*R100K/E107V*^ mutants, it is unclear whether the observed changes in *Pf*Actin quantity, ubiquitination, some other aspect of interactions with *Pf*Coronin, or a combination thereof, is most important in altering *Pf*Actin dynamics and driving ART resistance. However, these changes in *Pf*Actin dynamics are underlined by the hypersensitivity of *Pfcoronin*^*R100K/E107V*^ mutant parasites to JAS treatment.

Despite the importance of endocytosis and vesicular trafficking for parasite growth, the proteins and processes involved in endocytosis in *P. falciparum* are still being elucidated, with a very limited repertoire of proteins implicated^5,29–32^. In addition to *Pf*Kelch13 and its interaction partners, these proteins include phosphoinositide-binding protein PX1^33^, VPS45^30^, and the host enzyme peroxiredoxin 6^31^. In model organisms, actin assembly, disassembly, and turnover generate the force for membrane invagination and vesicle scission^34,35^. This process is accompanied by a host of actin regulatory proteins that work in concert to regulate actin dynamics. Only a few canonical actin-regulating proteins are maintained in *P. falciparum*^36–39^; *Pf*Coronin is one such regulatory protein.

Several genes involved in ART resistance have been implicated in hemoglobin uptake^4,40^ and digestion^41^. The most-studied ART resistance gene is *Pfkelch13*^4^; mutations in *Pfkelch13* confer ART resistance by reducing *Pf*Kelch13 protein abundance. *Pfkelch13* is localized to the cytostome neck and interacts with various proteins, including *Pf*UBP1 and AP2-μ that were implicated in ART resistance in separate studies^5,40^. Disruption of *Pfkelch13* or its interaction partners, in most cases, causes both defects in endocytosis (of RBC contents) and resistance to ART^5^. The endocytic and vesicular trafficking pathways in *P. falciparum* are not yet well-characterized, but some differences from the mammalian pathways have already surfaced. Notably, the cytostome appears to be devoid of clathrin, which mediates the most well-understood pathway for endocytosis in model organisms. On the other hand, AP2-μ, which typically works as part of the clathrin system, was observed^5^. Distinctions between mammalian and parasite processes may provide opportunities for novel drug-targeting strategies.

Prior studies on *Pf*Kelch13 showed that it is localized to the neck of the cytostome and facilitates hemoglobin uptake. The current study shows that *Pf*Coronin also facilitates hemoglobin uptake, but is localized to other structures, including the PPM/PVM, DV membrane, additional small vehicles, and preDVs (Fig. 6). Coronin lacks any obvious post-translational lipidation motifs or a transmembrane domain, but likely associates with membranes through interaction with phosphatidylinositol 4,5-bisphosphate (PI(4,5)P2)^42^. Associations between *Pf*Kelch13 and *Pf*Coronin were not detected in this study or in a prior study on *Pf*Kelch13^5^. It therefore seems likely that *Pf*Kelch13 and *Pf*Coronin contribute to separate steps of the same endocytosis process, or function in parallel endocytic processes. These possibilities are consistent with our previous *in vitro* genetic studies, which demonstrated epistasis between *Pfcoronin*^R100K/E107V^ and *Pfkelch13*^C580Y^ mutations in the Pikine genetic background^9^.

**Fig. 6:**
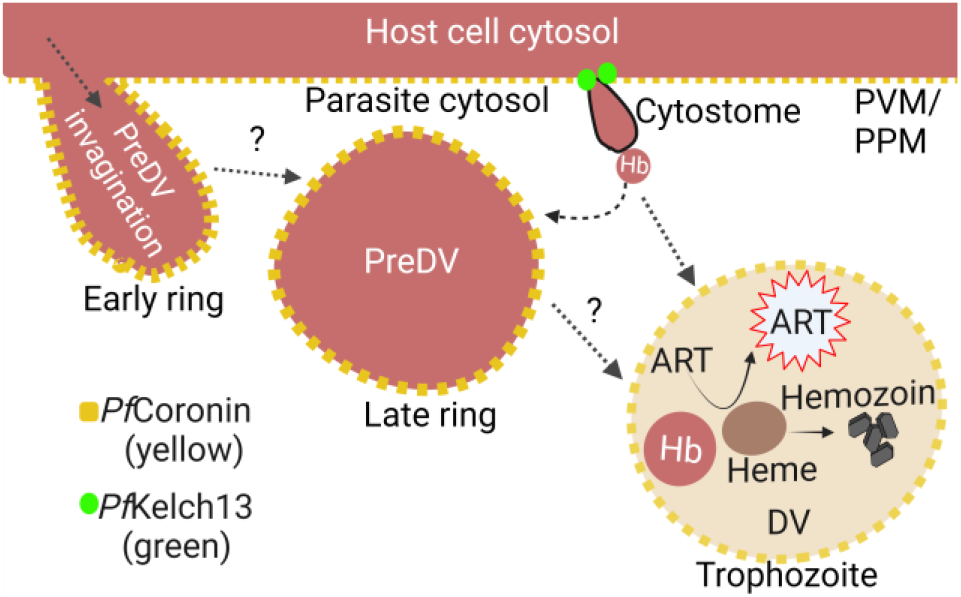
Working model for DV development, endocytosis, and *Pfcoronin*-mediated ART resistance. Diagram of infected RBC showing our current model. *Pf*Coronin is required for normal development of endocytic structures, such as the preDV. Mutations in *Pfcoronin* disrupt endocytosis and reduce uptake of hemoglobin from the host cell. This effect is similar to what’s seen for *Pf*Kelch13, but may occur via a different pathway. The theory is that after hemoglobin is digested in the DV, the free heme activates ART. Therefore, reduced endocytosis—via dysregulation of *Pf*Coronin or *Pf*Kelch13--decreases the amount of activated ART.

There is a massive gap in knowledge regarding the uptake and processing of hemoglobin, as well as the formation of the DV. DV formation is believed to rely on *de novo* biogenesis after invasion, and the DV is not visible in early stages^23,25,43,44^. A previous study in *P. falciparum*^23^ first identified a large hemoglobin-containing vacuole during the initial ring stage^23^. Another study^44^, using transmission electron tomography, described a somewhat similar large vacuole-like structure, but said that it remains connected to the RBC (i.e. an invagination) throughout the entire development of the ring stage (12-24 hpi). However, a separate transmission electron tomography of the ring stage of *P. chabaudi* observed both a single large hemoglobin-containing structure, entirely contained within the parasite, and multiple small hemozoin-filled vacuoles in ring-stage parasites. This study suggested that hemoglobin uptake in the ring stage parasites is mediated via both structures, while hemoglobin uptake is completely cytostome-mediated in later trophozoite stages^45^.

Recent advances in microscopy, such as U-ExM, simultaneously preserve molecular details and improve spatial resolution, allowing for a more detailed examination of ultrastructures in *P. falciparum*^20,21^. This technique is particularly advantageous in examinations of early rings, which are very small. Our U-ExM studies highlighted a large *Pf*Coronin-labeled vacuole, or preDV, which is entirely contained within the parasite in older rings (> 9hpi), while early rings (< 9hpi) have invagination structures connected to the RBC cytosol. We did not observe additional smaller vesicles at this stage but cannot rule out the possibility that these vesicles may exist, since small vesicles could be difficult to observe using U-ExM, especially if they were not *Pf*Coronin-coated. However, we do observe small *Pf*Coronin-coated vesicles in trophozoites and schizonts.

The possibility of multiple hemoglobin uptake pathways, via small vs. large vesicles, as described by Wendt et al^45^. is intriguing. This possibility would be consistent with our theory that *Pfcoronin* and *Pfkelch13* regulate two separate uptake processes—only the single large vacuole is *Pf*Coronin-coated in early rings. PreDV morphology is perturbed in *Pfcoronin* mutant parasites, suggesting that *Pf*Coronin—likely via *Pf*Actin regulation—is involved in formation of this structure. Further studies are needed to precisely delineate the process of DV formation and hemoglobin uptake in the ring-stage parasites, as well as the specific roles of *Pf*Coronin and *Pf*Kelch13.

ART remains a crucial component of front-line antimalarial therapies, and its antimalarial efficacy must be protected. The current study underlines the potential threat of *Pfcoronin*-mediated resistance—in fact, the *Pfcoronin*^*R100K*^ mutation was recently reported in Kenya, although artemisinin sensitivity/resistance was not assessed^46^. Other *Pfcoronin* polymorphisms were recently identified in Uganda^47^, and these polymorphisms were associated with ART resistance. We expect that surveillance of the *Pfcoronin* locus, in addition to *Pfkelch13*, will help protect malaria control efforts and guide decision-making to curtail the spread of drug resistance in Africa. Additionally, we hope that mechanistic understanding of *Pfcoronin*-mediated ART resistance will inform strategies to defend the efficacy of ART therapies.

## Acknowledgments

Funding: NIH (5R01AI099105-07) to DFW and DLH and NIH (R01 AI145941) to JDD.

## Author contributions

Conceptualization: I.U., J.D.D., D.L.H., S.K.V., S.B., S.A., and D.F.W.; Methodology: I.U., M.A.F., A.Y.B., E.H., B.C.W., M.K., S.H.S., A.I.S., K.L.S., M.C.M., S.A.,; Data analyses: I.U., A.Y.B., B.C.W., M.K., M.C.M, S.H.S., S.A.,; Writing - original draft: I.U.; writing - review & editing: I.U., J.D.D., D.L.H., S.K.V., S.B., S.A., and D.F.W.; Funding acquisition: D.F.W., and D.L.H., Supervision: D.F.W.

