## Supplementary_Figures for "Artemisinin resistance mutations in *Pfcoronin* impede hemoglobin uptake"

*Corresponding authors:

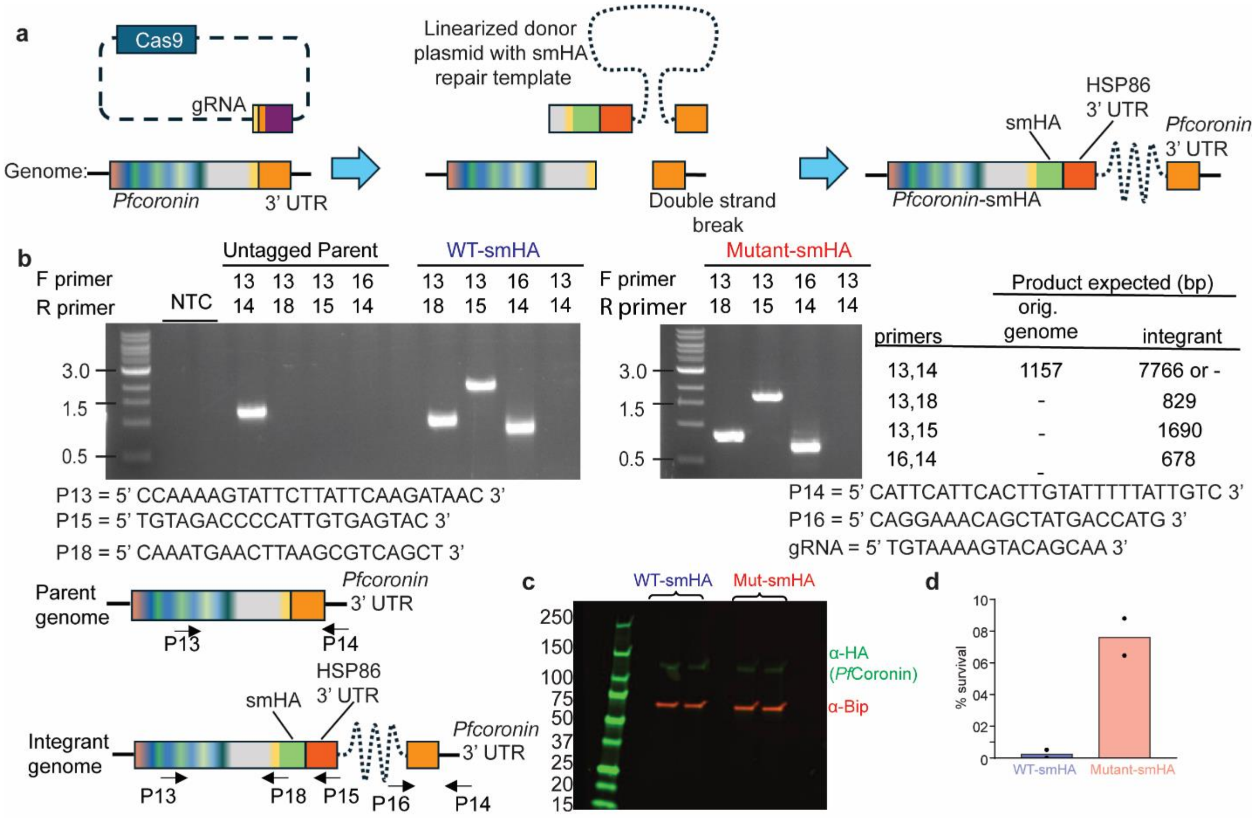


**Supplementary Fig. 1:** Generation of spaghetti monster HA tag (smHA)-tagged *Pfcoronin*$^{WT}$-*smHA* and *Pfcoronin*$^{R100K/E107V}$-*smHA* transgenic lines. (a) Schematic for C-terminal endogenous tagging of WT (*Pfcoronin*$^{WT}$) and mutant (*Pfcoronin*$^{R100K/E107V}$) parasites with smHA. Cas9 created a double-stranded break near the C-terminus of the *Pfcoronin* locus in either the *Pfcoronin*$^{WT}$ or the *Pfcoronin*$^{R100K/E107V}$ parent strain. This break was subsequently repaired via double crossover/homologous recombination between the *Pfcoronin* coding sequence (CDS; upstream of the break), the *Pfcoronin* 3' UTR, and matching sequences on the donor vector. This resulted in the integration of the entire donor vector, including an smHA tag appended to the C-terminus of *Pfcoronin.* (b) PCR confirmation of smHA tagging in *Pfcoronin*$^{WT}$-*smHA* and *Pfcoronin*$^{R100K/E107V}$-*smHA* lines. Expected sizes from PCRs using genomic DNA templates from untagged (parental/ "orig. genome") *Pfcoronin* and *Pfcoronin-smHA* ("integrant") lines are detailed in the associated table. Primers used for the PCRs are shown and their binding positions are annotated in the schematic of the parasite genome. The gRNA sequence used is also shown. (c) Confirmation of smHA-tagged *Pfcoronin*$^{WT}$-*smHA* and *Pfcoronin*$^{R100K/E107V}$-*smHA* lines via western blotting using anti-HA tag (mouse mAb 2-2.2.14; green) and anti-BiP (loading control; rabbit polyclonal red) antibodies; samples were generated from late-stage parasites. Two biological replicates are shown for each strain. (d) Tightly synchronized 3-hour-old ring-stage *Pfcoronin*$^{WT}$-*smHA* and *Pfcoronin*$^{R100K/E107V}$-*smHA* parasites were pulsed with 700 nM DHA for 6-hours, and survival was assessed 66 hours later. As expected, substantial survival (i.e., artemisinin resistance) was observed in the *Pfcoronin*$^{R100K/E107V}$-*smHA* line. Generally, the minimum survival threshold for ART resistance is 1%. Means from two independent biological replicates are shown.

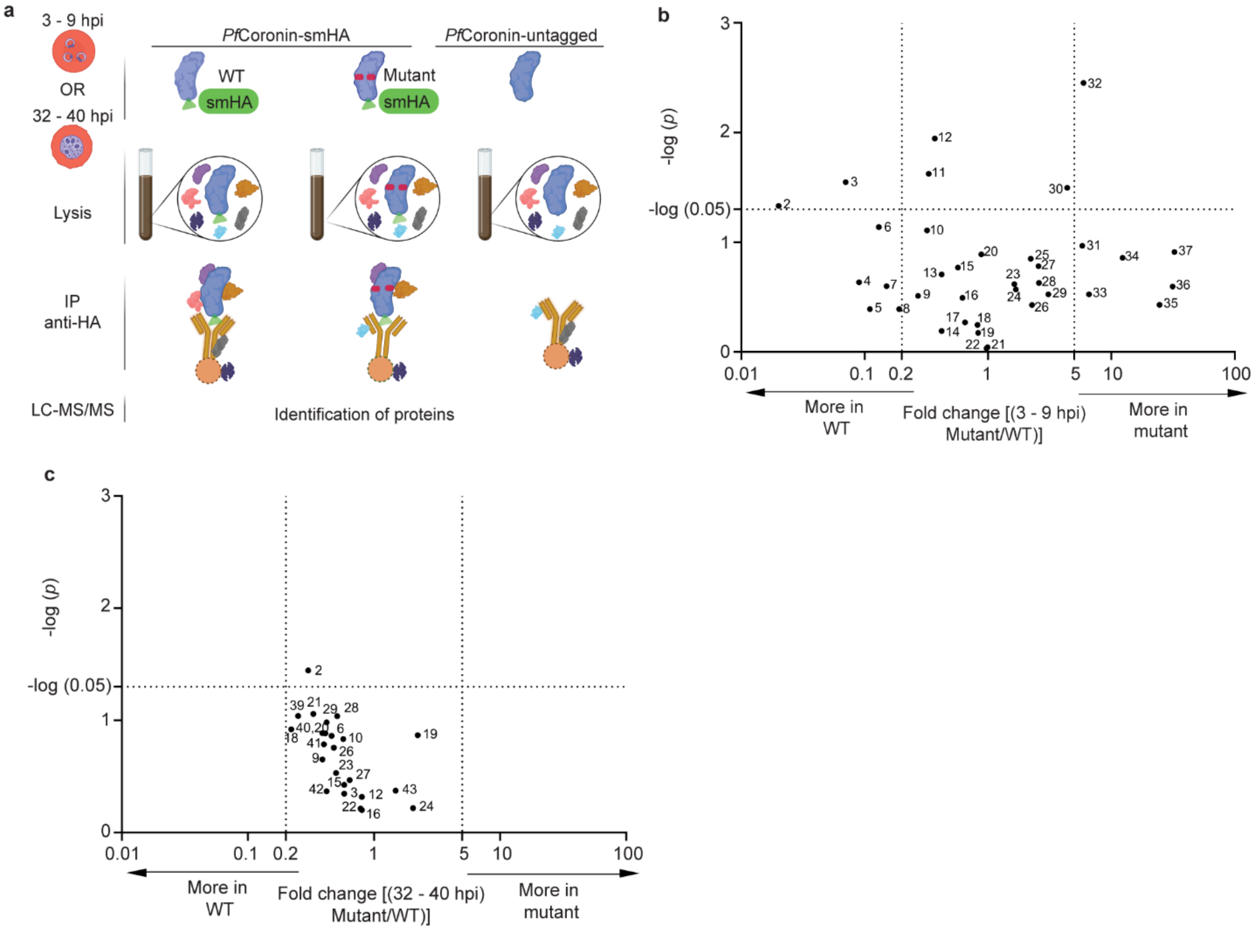


**Supplementary Fig. 2:** (a) Immunoprecipitation of *Pf*Coronin-smHA and associated proteins from young ring-stage (3-9 h) and late-stage (32-40 h) parasites using anti-HA antibodies. Co-purified proteins were identified by LC-MS/MS, with an untagged *Pfcoronin*$^{WT}$ control used to filter nonspecific interactions. hpi = hours post invasion. (b-c) A volcano plot of hit interaction partners shows changes between *Pf*Coronin$^{WT}$-smHA and *Pf*Coronin$^{R100K/E107V}$-smHA in (b) rings, (c) trophozoites. The x-axis shows the fold change between *Pf*Coronin$^{WT}$-smHA and *Pf*Coronin$^{R100K/E107V}$-smHA bait proteins. The Y axis shows the statistical significance of the change between *Pf*Coronin$^{WT}$ and *Pf*Coronin$^{R100K/E107V}$ (unpaired t-test). The 37 hit proteins (for rings in b) and 24 hit proteins (for trophozoites in c) plotted on the volcano plot were found in all bioreps (either *Pf*Coronin$^{WT}$-smHA or *Pf*Coronin$^{R100K/E107V}$-smHA) and have at least 5% of the *Pf*Coronin protein abundance. In rings (b) two proteins (#1, *Pf*3D7_0517700, eukaryotic translation initiation factor 3 subunit B; and #38, *Pf*3D7_1424400, 60S ribosomal protein L7-3, putative) fulfilled these criteria but could not be displayed because they were detected only in *Pf*Coronin$^{WT}$-smHA (*Pf*3D7_0517700) or only in *Pf*Coronin$^{R100K/E107V}$-smHA pulldowns (*PF*3D7_1424400), so a log fold change could not be computed. A full list of hit proteins can be found in Supplementary Data 3 (for rings) and Supplementary Data 4 (for trophozoites). Proteins of interest are defined as either those with $p < 0.05$ (above the dotted line on the y-axis), or a 5-fold or greater change between *Pf*Coronin$^{WT}$-smHA and *Pf*Coronin$^{R100K/E107V}$-smHA parasites (to the left (0.2) or right (5) of the vertical dotted lines on the x-axis). The proteins are numbered and are detailed in Supplementary Data 3 (for rings) and Supplementary Data 4 (for trophozoites).

Each data point represents two or three independent biological replicates for rings and trophozoites, respectively.

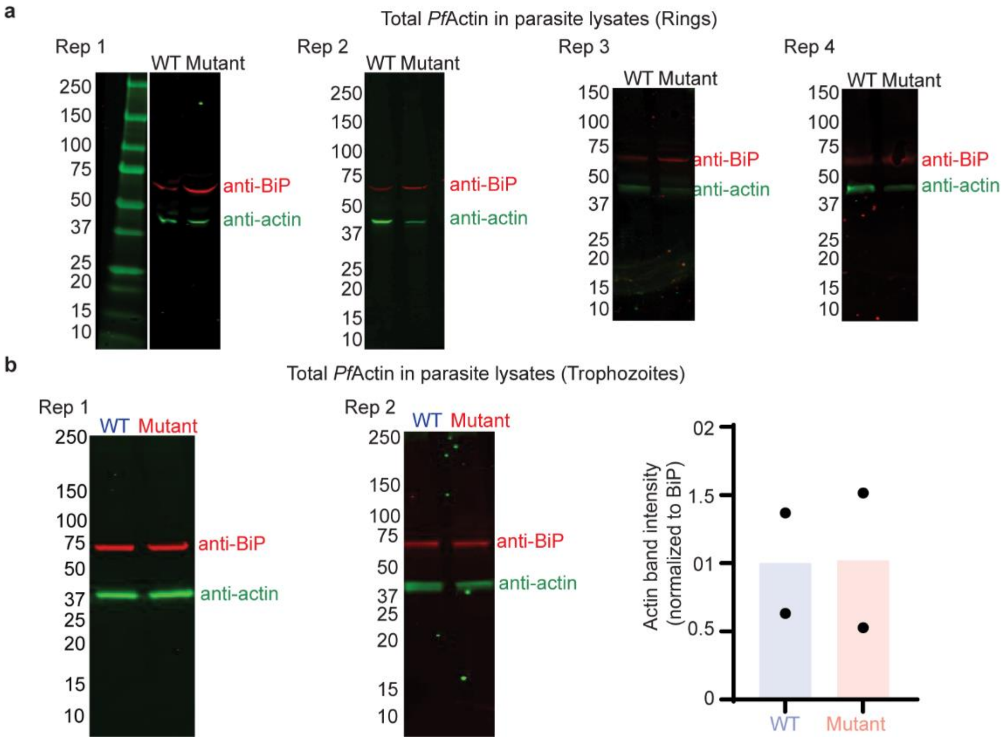


**Supplementary Fig. 3:** (a) Total *Pf*Actin is reduced in parasites expressing *Pf*Coronin$^{R100K/E107V}$-smHA (mutant), compared to *Pf*Coronin$^{WT}$-smHA (WT). Lysates from parasites expressing *Pf*Coronin$^{R100K/E107V}$-smHA were compared by western blot to lysates from parasites expressing *Pf*Coronin$^{WT}$-smHA. Blots were probed with anti-actin (mouse monoclonal; green) and anti-BiP (loading control; rabbit polyclonal, red) antibodies. All western blots quantified in Fig. 1d (biological replicates 1-4) are shown. (b) Total *Pf*Actin in parasites expressing *Pf*Coronin$^{R100K/E107V}$-smHA (mutant), compared to *Pf*Coronin$^{WT}$-smHA (WT) is not changed in late trophozoite stage parasites. Blots were probed with anti-actin (mouse monoclonal; green) and anti-BiP (loading control; rabbit polyclonal, red) antibodies. Quantification was based on the two independent biological replicates shown.

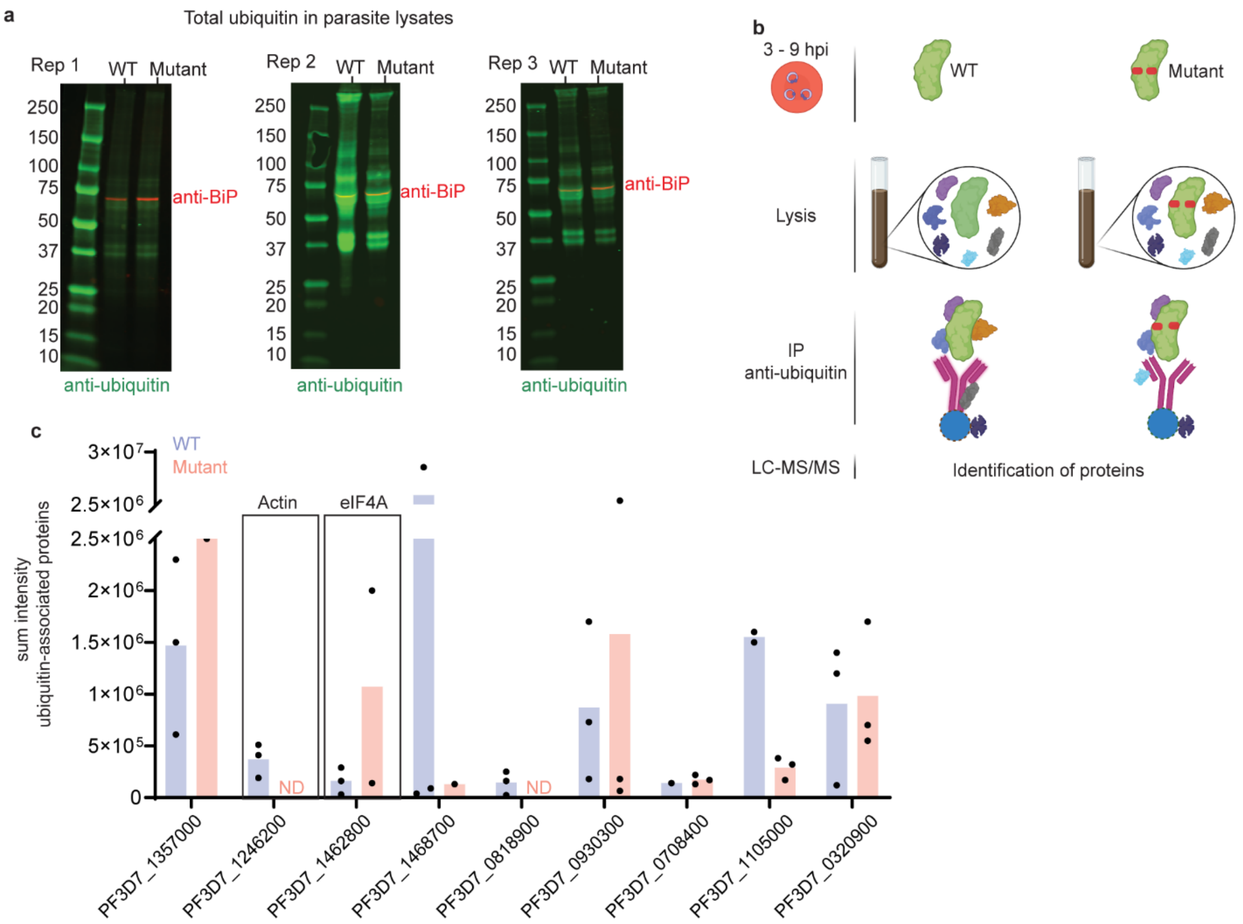


**Supplementary Fig. 4:** (a) No gross differences in global ubiquitination were observed in parasites expressing *Pf*Coronin$^{R100K/E107V}$-smHA (mutant), compared to *Pf*Coronin$^{WT}$-smHA (WT). Lysates from parasites expressing *Pf*Coronin$^{WT}$-smHA and *Pf*Coronin$^{R100K/E107V}$-smHA were compared by western blot. Blots were probed with anti-ubiquitin (mouse mAb FK2, green) and anti-BiP (loading control; rabbit polyclonal red) antibodies. Three independent biological replicates are shown. (b) Schematic showing immunoprecipitation of ubiquitin and associated proteins from young ring-stage parasites (3-9 h). (c) Comparison of *Pf*Ub-associated hit protein abundances from parasites expressing *Pf*Coronin$^{WT}$-smHA vs *Pf*Coronin$^{R100K/E107V}$-smHA. Following anti-ubiquitin IP, co-purified proteins were identified and quantified by LC-MS/MS. Proteins shown were identified in all replicates of either *Pf*Coronin$^{WT}$-smHA or *Pf*Coronin$^{R100K/E107V}$-smHA IPs. *Pf*Actin and *Pf*eIF4A, the only two proteins that were also identified as *Pf*Coronin interaction partners (anti-HA IPs), are highlighted with boxes. Data shown are average measurements from three independent biological replicates. Complete results are shown in Supplementary Data 5.

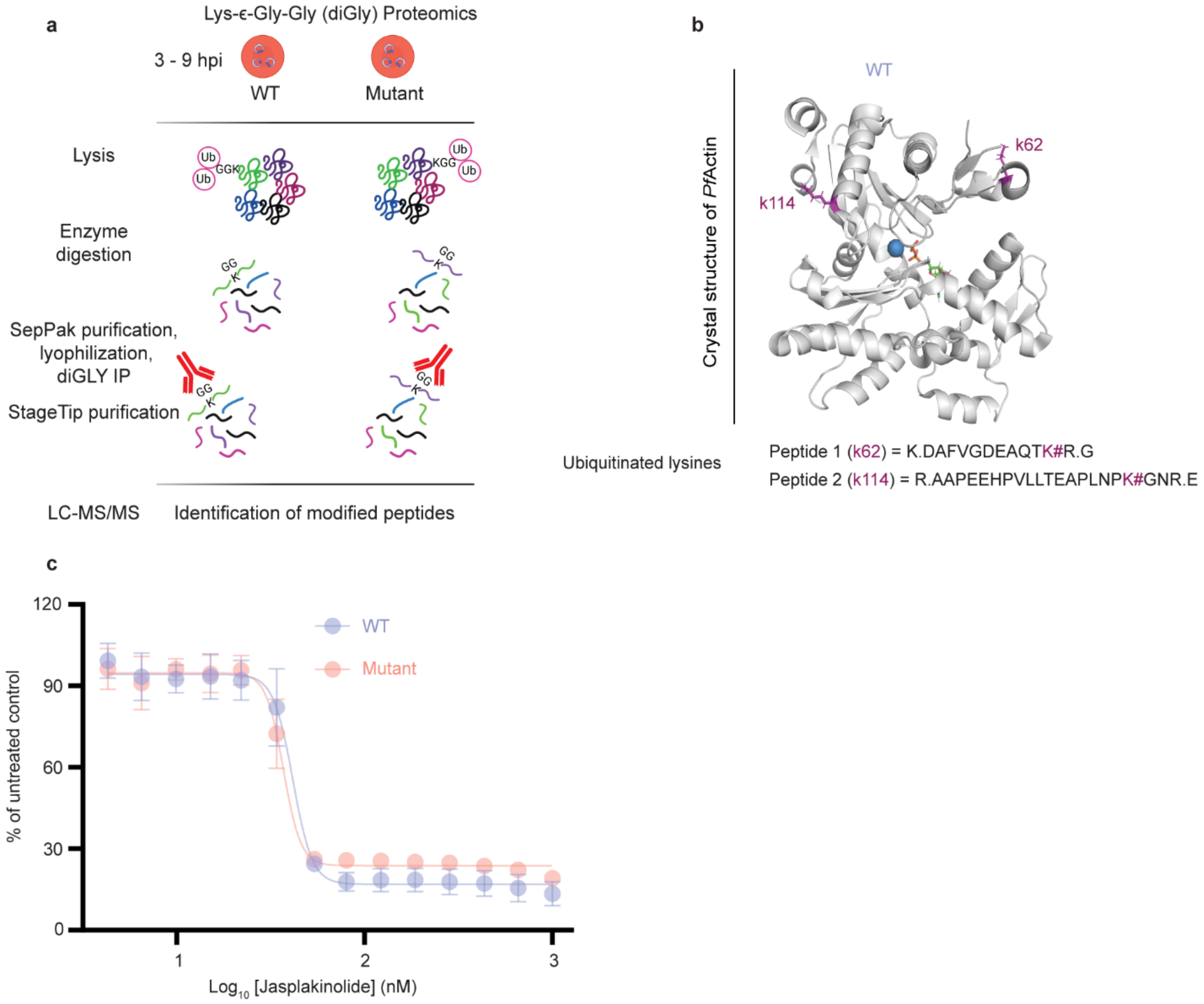


**Supplementary Fig. 5:** Lys-ϵ-Gly-Gly (diGly) proteomics to identify proteins that are ubiquitinated at specific sites. (a) Young ring-stage *Pfcoronin*$^{WT}$ and *Pfcoronin*$^{R100K/E107V}$ parasites (3-9 h) were lysed, normalized by protein concentration, and subjected to trypsin digestion. Peptides were enriched with an anti-diGly antibody, and peptides were identified by LC-MS/MS. (b) Crystal structure of *Pf*Actin[1] . The ubiquitination sites identified are labelled and the individual peptides identified are detailed in Supplementary Data 6. (c) A standard 72-h assay shows no difference between WT ($EC_{50}$ = 40.1 ± 1.3 nM) and mutant ($EC_{50}$ = 37 ± 2.6 nM). Data points shown are average measurements from three independent biological replicates.

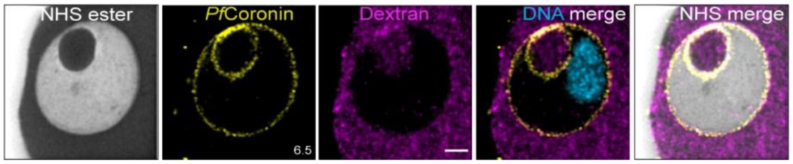


**Supplementary Fig. 6:** A representative image from ultrastructure expansion microscopy (U-ExM), showing *Pf*Coronin-labeled intracellular putative pre-DV compartment (PPDC) housing host cell contents, as indicated by dextran staining. This pattern is consistent across all analyzed ring-stage PPDCs in *Pfcoronin*$^{WT}$ parasites. Alexa Fluor 405-conjugated NHS ester (protein density), HA (*Pf*Coronin, yellow), Streptavidin 488 (dextran, magenta), Sytox Deep Red (DNA, Cyan). The images are maximum-intensity projections, with the number indicating the Z-axis thickness in µm. Scale bars: 2 µm

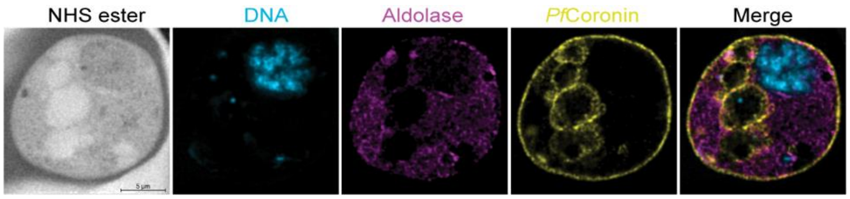


**Supplementary Fig. 7:** A representative image from ultrastructure expansion microscopy, showing *Pf*Coronin-labeled intracellular chains of small and large vacuoles in a trophozoite. Alexa Fluor 405-conjugated NHS ester (protein density), Sytox Deep Red (DNA, cyan), Aldolase 555 for parasite cytoplasm (magenta), HA (*Pf*Coronin, yellow). The images are maximum-intensity projections.

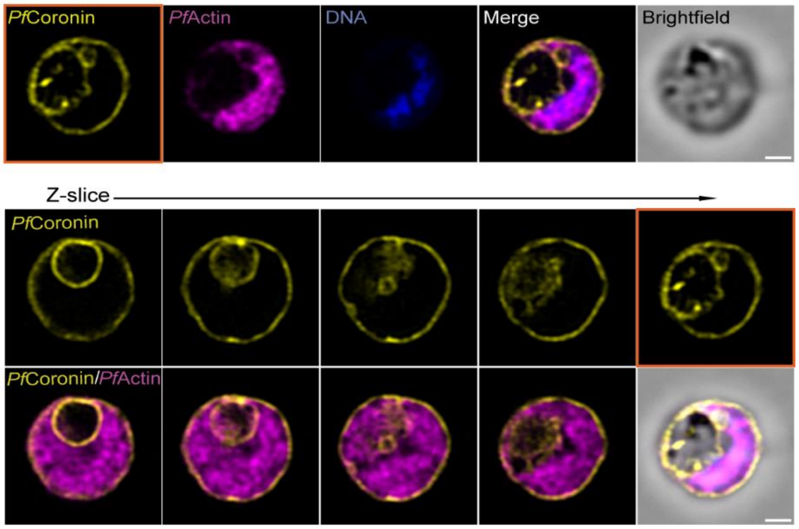


**Supplementary Fig. 8:** Super-resolution fluorescence microscopy using Zeiss LSM980 with Airyscan2 shows localization of *Pf*Coronin and *Pf*Actin early schizont stage parasites. Scale bar = 2 µm.

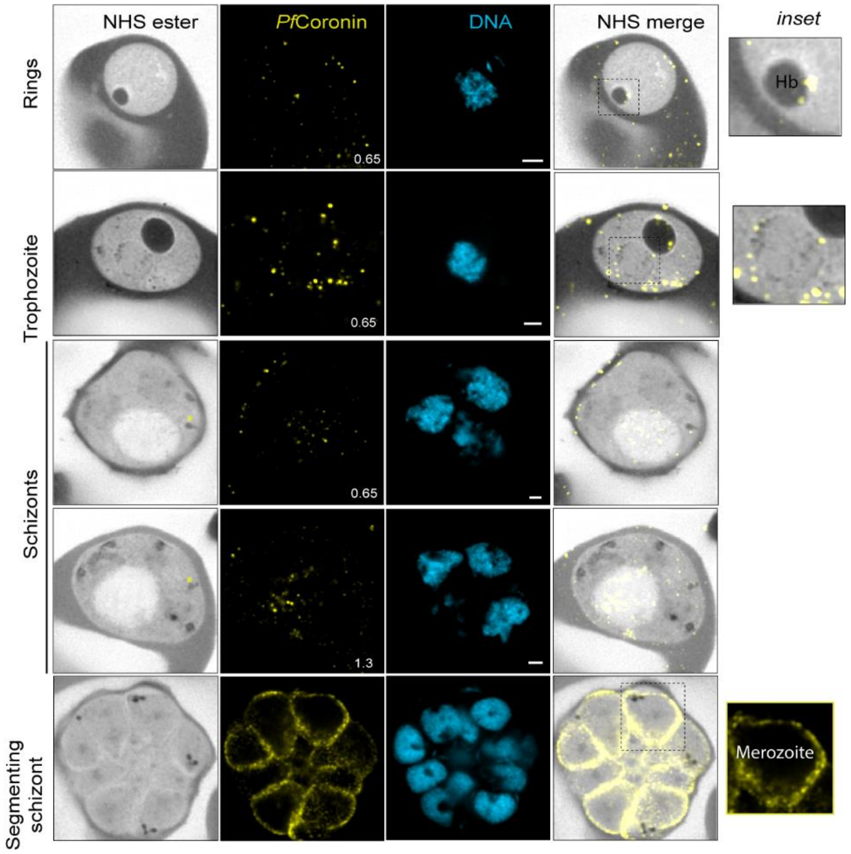


**Supplementary Fig. 9:** U-ExM shows *Pf*Coronin localization throughout the blood-stage cycle in *Pfcoronin*$^{R100K/E107V}$*-smHA* mutant parasites. Cytostomes are marked with a yellow asterisk in the first (NHS ester) panel of each row. Alexa Fluor 405-conjugated NHS ester (protein density), HA (*Pf*Coronin, yellow), Sytox Deep Red (DNA, cyan). Images are maximum-intensity projections, number on image = Z-axis thickness of projection in µm. Scale bars = 2 µm.

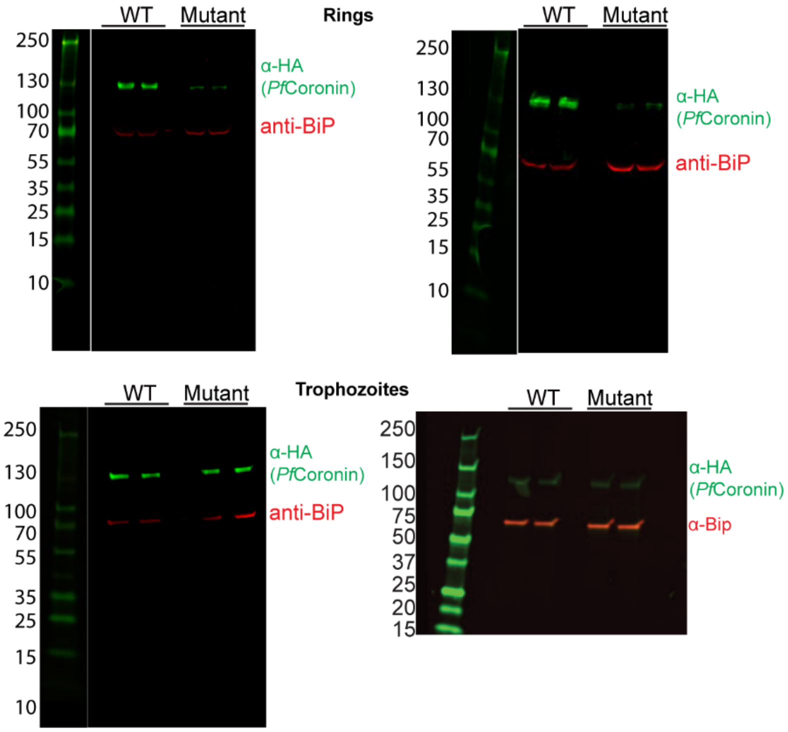


**Supplementary Fig. 10**: Total *Pf*Coronin levels in lysates from parasites expressing *Pf*Coronin$^{R00K/E107V}$-smHA (mutant) were compared by western blot to lysates from parasites expressing *Pf*Coronin$^{WT}$-smHA. Blots were probed with anti-HA (mouse monoclonal; green) and anti-BiP (loading control; rabbit polyclonal, red) antibodies. All western blots quantified in Fig. 4b are shown.

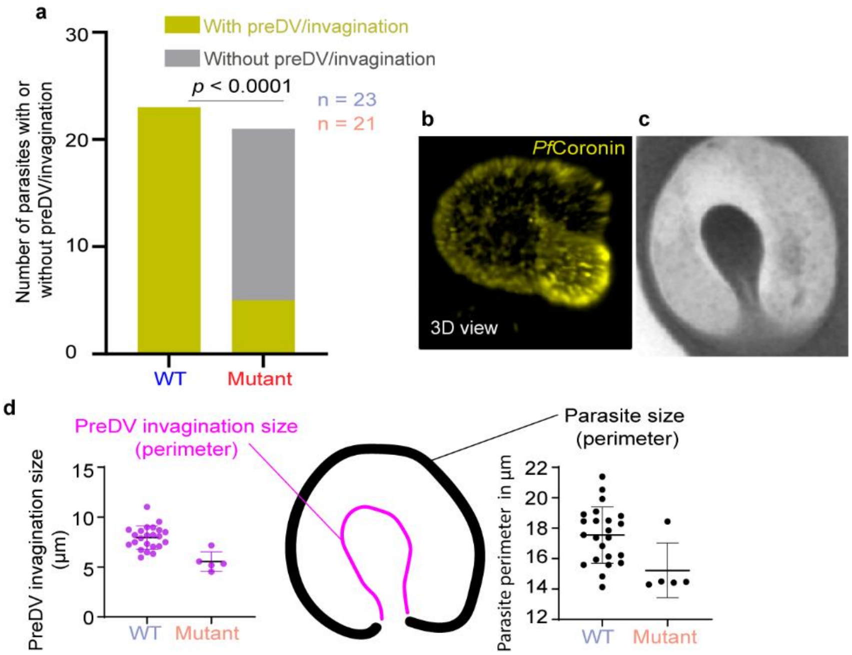


**Supplementary Fig. 11:** (a) *Pfcoronin*$^{WT}$*-smHA* and *Pfcoronin*$^{R100K/E107V}$*-smHA* (mutant) parasites at the early ring stage (< 9hpi), compared for the presence of a preDV/invagination, analyzed using Fisher's exact test (Baptista-Pike method). (b) 3D view of *Pf*Coronin staining (anti-HA) of a ring-stage parasite, displaying the parasite body and preDV invagination. (c) NHS-ester staining shows the parasite body and invagination. (d) Cartoon showing the measurement of preDV/invagination (left graph in magenta showing individual measurements) and parasite size (right graph showing individual measurements of parasite size). PreDV/invagination sizes (left graph) for both WT and mutant parasites were normalized against parasite size (right graph), and the resulting data are presented in Fig 5b.

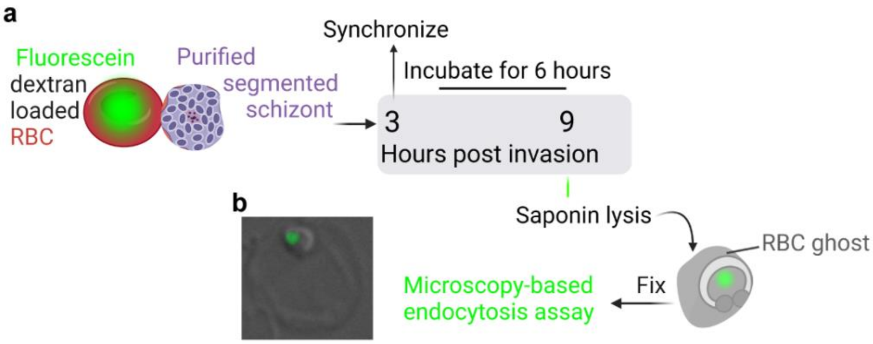


**Supplementary Fig. 12:** (**a**) Diagram of sample preparation for endocytosis. RBCs were lysed in a hypotonic solution containing fluorescein-dextran. After re-sealing the RBCs (trapping the fluorescein dextran within), schizonts were added and allowed to re-invade for 3 hours. As the parasites (rings) grew within the fluorescein-dextran-containing RBCs, fluorescein-dextran was endocytosed and accumulated in the DV and other compartments. The parasites were then synchronized to remove any remaining schizonts and allowed to grow for a further 6 hrs, closely mimicking the RSA setup. RBC cell contents were removed by saponin lysis and parasites were fixed for microscopy. (**b**) DIC and AF488 channel images were captured using a 100X oil immersion objective on a Zeiss Axio Observer. Representative image is shown. Images were blinded prior to analysis. Analysis was carried out for regions of interest (ROIs) where an RBC ghost could be visualized around a green channel signal in the merged image. Vesicle size (area), internalized fluorescence intensity (mean gray value), and integrated density (over the entire vesicle) of green-channel fluorescence signal was assessed.

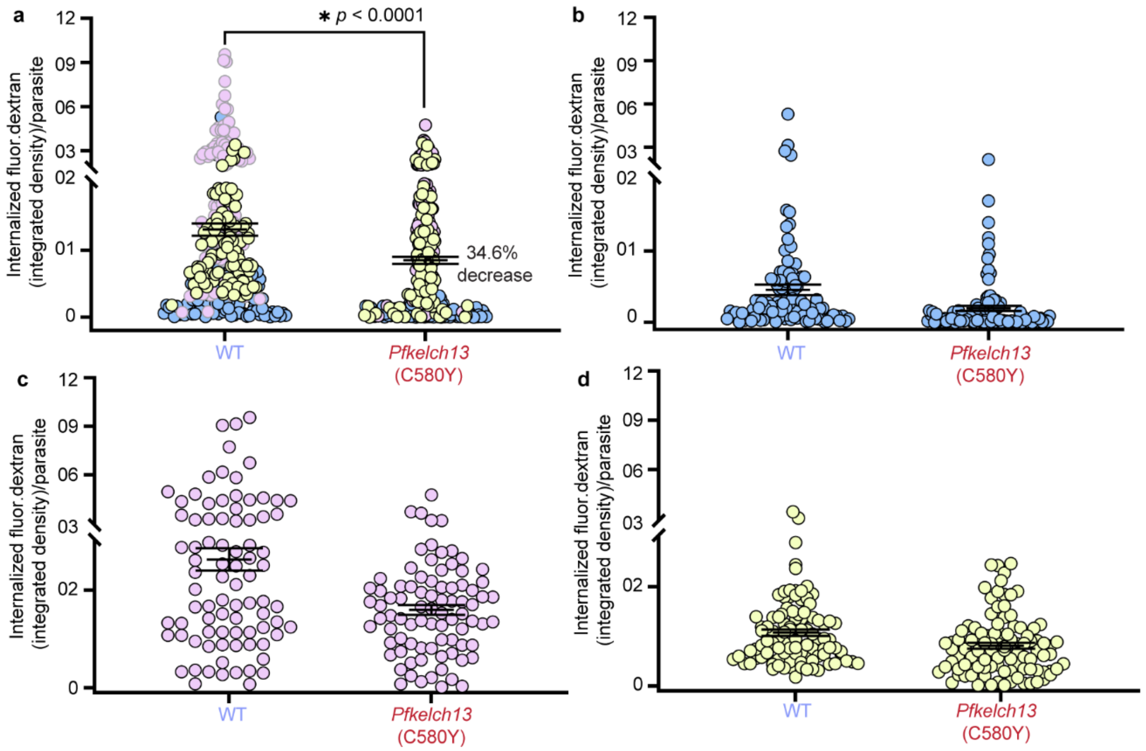


**Supplementary Fig. 13:** Comparison of internalized fluorescein-dextran in WT vs. *Pfkelch13* (C580Y) parasites by (a) integrated density (vesicle size x intensity). WT *Pfcoronin* n = 286; *Pfkelch13* (C580Y) n = 286. Each data point represents fluorescein-dextran uptake by a single parasite. *p*-values (unpaired, nonparametric Mann-Whitney test) and % reduction of the mean is indicated. These data were pooled from the individual biological replicates shown in b-d.

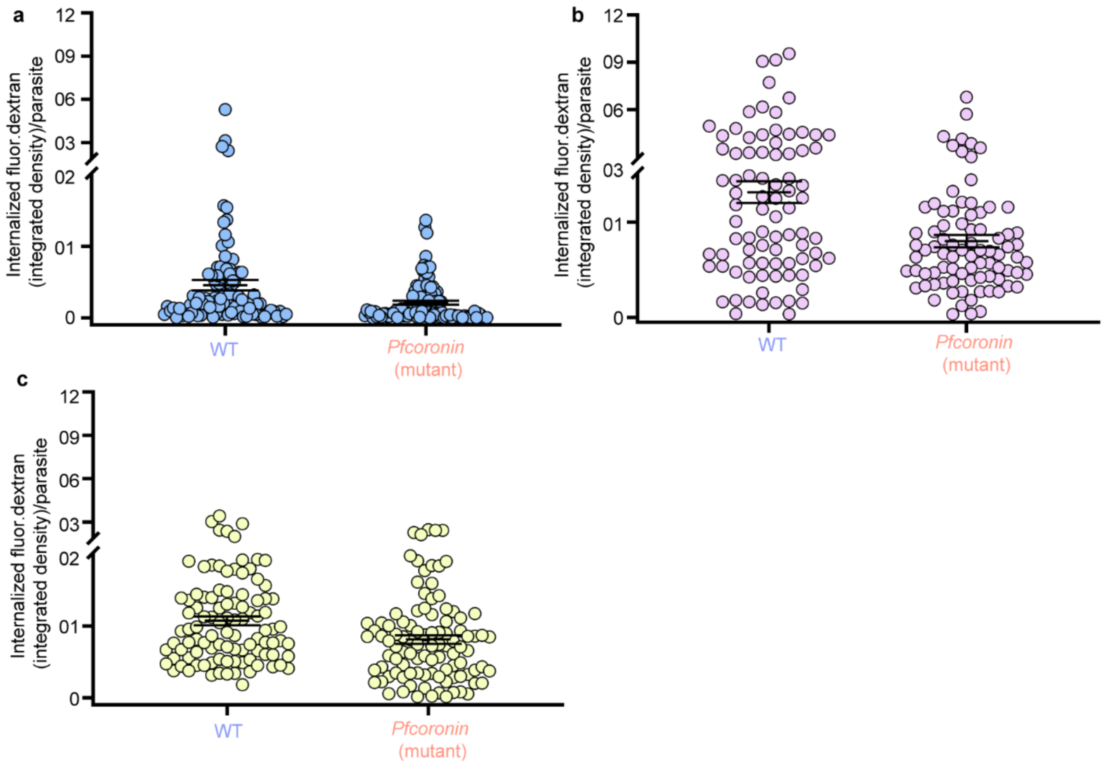


**Supplementary Fig. 14:** (a-c) Individual biological replicates of internalized fluorescein-dextran in WT vs. *Pfcoronin* (R100K/E107V) mutant ring-stage parasites (3-9 hours post-invasion) quantified and pooled in Fig. 5. Each data point represents fluorescein-dextran uptake by a single parasite.
