## Supplementary material for "Artemisinin resistance mutations in *Pfcoronin* impede hemoglobin uptake": Methods

**Parasite culture:** *P. falciparum* culture-adapted field isolates, Pikine (SenP019.04), Pikine_R (DHA selected line[1], used for Jasplakinolide RSAs), CRISPR-engineered Pikine *Pfcoronin*$^{R100K/E107V}$ mutant (CRISPR-edited, used for timecourse experiments), *Pf*kelch13$^{C580Y}$ mutant (CRISPR-edited, used for endocytosis)[1,2], spaghetti monster HA tag (smHA)-tagged Pikine *Pfcoronin*$^{WT}$*-smHA*, and Pikine *Pfcoronin*$^{R100K/E107V}$*-smHA* mutant lines (used for all experiments unless specified otherwise) were maintained in complete RPMI 1640 medium (Gibco) supplemented with 10% $O^+$ serum at 4% hematocrit (unless specified otherwise) in an atmosphere of 1% O2, 5% CO2, and 94% N2 at 37 °C[1,2]. All smHA-tagged lines were additionally cultured with 5nM WR99210 (Jacobus Pharmaceutical). Staging and parasitemia were assessed by light microscopy of Giemsa-stained thin blood smears[3,4]. The parasites were synchronized using sequential sorbitol lysis treatment [5].

**Generation of smHA-tagged parasites and transfection**: *Pfcoronin* (PF3D7_1251200) was tagged with smHA using CRISPR/Cas9. The CRISPR guide sequence (selected using benchling.com) was 5'-AATGTGTAAAAGTACAGCAA-3', which matches a site near the C-terminus of the *Pf*Coronin coding sequence. A double-stranded DNA segment with "sticky ends" was generated by annealing oligonucleotides 5'-ATTGAATGTGTAAAAGTACAGCAA-3' and 5'-AAACTTGCTGTACTTTTACACATT-3'. This segment was ligated within the U6 cassette of pBAM203[6] (pre-digested with BbsI; kind gift of Dr. Jeffrey Dvorin, Harvard Medical School), proximal to the invariable part of the chimeric sgRNA scaffold, to generate pUF1-Cas9-hDHODH-Coronin. The donor/homologous repair plasmid for smHA-tagging *Pf*Coronin was derived from pSAB55[7] (kind gift of Dr. Jeffrey Dvorin, Harvard Medical School), which contains coding sequence for spaghetti monster-HA tag (smHA), followed by the *Pf*HSP86 3' UTR. Additionally, the plasmid includes a cassette for expression of human dihydrofolate reductase (hDHFR), allowing selection using WR99210. A 5' homology arm (plus shield mutations) was amplified from the end (C-terminus) of the *Pf*Coronin coding sequence using primers: 5'-CAATTGCCATGGGACAGTTTACCAAAAAATTTACCTTC-3' and 5'-ATCTCGAGTAAAACGGTTGCAGTACTTTTACACATTTTCATGTCTAG-3'; the second primer includes shield mutations to protect from cutting by Cas9. This segment was inserted into the NcoI/XhoI sites of pSAB55. A 3' homology arm was amplified from the *Pf*Coronin 3' UTR using 5'-CGTCAGGATCATCGCGGCCGCGGCACATCAGCGATCAC-3' and 5'-CATGGCAATTGAGATCTCCGAAAAATTTGTATGATATTGAAAAG-3'; this segment was inserted into the NotI/BglII sites of the plasmid using Gibson cloning, generating pSAB55-Coronin-smHA. Prior to transfection, the complete pSAB55-Coronin-smHA plasmid was linearized by digestion with NcoI. Plasmids (50 µg each of pUF1-Cas9-hDHODH-Coronin and linearized pSAB55-Coronin-smHA) were transfected into Pikine parasites carrying either *Pf*Coronin$^{WT}$ (WT) or *Pf*Coronin$^{R100K/E107V}$, as reported previously[2]. Transfections were performed as described previously[2,8]. Transfected parasites were allowed to recover overnight before the addition of 5 nM WR99210 (selection agent for pSAB55-Coronin-smHA) and 500 nM DSM1 (selection agent for pUF1-Cas9-hDHODH-Coronin). Drug selection with DSM1 was continued for 3 days. Parasites grew up around 3 weeks and were cloned by limiting dilutions prior to studies. Confirmation of the smHA-tagged parasites is detailed in Supplementary Fig. 1.

**Ring stage survival assay (RSA):** RSA was performed as described previously[1,2,9,10]. *P. falciparum* cultures at the late segmented schizont stage were purified using the Percoll gradient and allowed to reinvade for 3 hours before subjecting them to synchronization. Highly synchronized (0-3 hours post-invasion) parasites were pulsed with 700 nM DHA (Sigma-Aldrich) or 740 nM jasplakinolide (ref no, 420127, Sigma-Aldrich), or DMSO control. The pulse was followed by drug washout and further incubation at 37˚C for 66 hours, and parasitemia was assessed by light microscopy of Giemsa-stained thin blood smears[1,2]. RSA survival rates in percentage were estimated by comparing the survival of viable parasites to the untreated DMSO control for the same parasite line. Smears were blinded, and 10,000 RBCs were counted for each condition. Unless otherwise specified, all experiments used at least three independent biological replicates and two technical replicates.

**Concentration–response assays:** All $EC_{50}$s were estimated using the standard 72 hour Malaria SYBR green I fluorescence assay[11,12]. Ring-stage parasites (40 µL, 1% hematocrit and 1% starting parasitemia, n = 3) were cultured for 72 hours at 37°C in a 384-well black clear-bottom plates containing 12-point serial dilutions of test compounds, 10 µM DHA kill-control and no drug growth-control. Lysis buffer containing 16% w/v saponin, 1.6% Triton X-100, 5 mM EDTA, 20 mM Tris-HCl, pH 7.4) and 1x SYBR Green I fluorescent dye (CAT. No. S7563, Invitrogen) was added, and the plates were incubated at room temperature in the dark for at least 6 hours prior to reading. The fluorescent signal was measured using SpectraMax M5 (Molecular Devices, Sunnyvale, CA) plate reader (excitation at 494 nm, emission at 530 nm). Experiments were carried out as three technical triplicates on the same plate, with three independent biological repeats of each plate performed. Controls in each biological replicate consisted of parasite culture with no drug added (100%) or 10 µM DHA kill-control (0%). The mean and SD of fluorescence data from three independent biological repeats were expressed as a proportion of the untreated control (100%) and calculated as follows: $100 \times [\mu(S) - \mu(-)/\mu(+) - \mu(-)]$, where $\mu(S)$, $\mu(+)$ and $\mu(-)$ represent the means for the sample in question and 100% and 0% controls, respectively as previously described[4,13]. The percentage growth was plotted against $\log_{10}$-transformed drug concentration and $EC_{50}$s were estimated using a non-linear regression (log(inhibitor) vs response-variable slope with four parameters) in GraphPad Prism (GraphPad Software, Inc., San Diego, CA, USA). Values from the three biological replicates were used to calculate the mean $EC_{50}$ values ± SE shown.

**Co-Immunoprecipitation**: For co-immunoprecipitation, (Co-IP) experiments, tightly synchronized (with 5% D-sorbitol[5] and density centrifugation with Percoll) parasites were used. A total of $4 \times 10^9$ (32 to 40 hpi, for late stages) or $4 \times 10^9$ (3-9 hpi, for ring stages) parasites were harvested by washing with PBS, and then the RBCs were lysed with 0.05% saponin in PBS in the presence of EDTA-free complete mini protease inhibitors (CAT. No. 4693159001, Sigma). The samples were then washed with PBS supplemented with protease inhibitors cocktail to remove residual RBC material. Parasite extracts were lysed in RIPA buffer (50 mM Tris-HCl, pH 7.5, 150 mM NaCl, 1% NP-40, 0.5% sodium deoxycholate, 0.1% sodium dodecyl sulfate, and EDTA-free complete mini protease inhibitors) by incubating the samples on a rotator for 1 hour at room temperature. The samples were then sonicated three times at 20% amplitude for 30 s at 4 °C with 3-minute rests on ice to chill the samples. The soluble fractions were collected by spinning the samples 12,000 g for 30 minutes and incubated with 40 µL anti-HA magnetic beads (ref no 88836, Pierce) overnight at 4 °C. For co-IP with anti-ubiquitin mAb FK2 antibody (ref no ST1200, sigma), the soluble fractions were incubated with the antibody overnight at 4 °C and then applied to protein A/G magnetic beads (CAT. No. 88802, Pierce) for 1 hour at room temperature, following the manufacturer's instructions. Beads were washed three times with RIPA in the presence of protease inhibitor cocktail, each time with a 10-minute incubation at RT on a rotator. The beads were subsequently washed once each with buffer 1 (2% SDS in $dH_2O$), buffer 2 (50 mM HEPES, pH 7.5, 500 mM NaCl, 1 mM EDTA, 1% Triton-X 100 and 0.1% sodium deoxycholate in $dH_2O$) and buffer 3 (10 mM Tris-HCl pH 8.0, 250 mM lithium chloride, 1 mM EDTA, 0.5% NP-40 and 0.5% sodium deoxycholate in $dH_2O$). The beads were then washed at least six times with buffer 4 (50 mM ammonium bicarbonate in $dH_20$), and resuspended in 40 µL of 5 mM ammonium bicarbonate. Then, 200 ng/µL of modified sequencing-grade trypsin (Promega, Madison, WI) was spiked in and the samples were incubated overnight at 37 °C. The samples were then placed on a magnetic plate, and the liquid was removed. The extracts were then dried in a speed-vac (~1 hour). Samples were then re-suspended in 50 µL of high-performance liquid chromatography (HPLC) solvent-A (2.5% acetonitrile, 0.1% formic acid) and desalted by STAGE tip[14]. On the day of analysis, the samples were reconstituted in 10 µL of HPLC solvent-A. A nano-scale reverse-phase HPLC capillary column was created by packing 2.6 µm C18 spherical silica beads into a fused silica capillary (100 µm inner diameter, ~30 cm length) with a flame-drawn tip[15]. After equilibrating the column, each sample was loaded via a Famos autosampler (LC Packings, San Francisco CA) onto the column. A gradient was formed, and peptides were eluted with increasing concentrations of solvent-B (97.5% acetonitrile, 0.1% formic acid). As peptides eluted, they were subjected to electrospray ionization and then entered into an LTQ Orbitrap Velos Elite ion-trap mass spectrometer (Thermo Fisher Scientific, Waltham, MA). Peptides were detected, isolated, and fragmented to produce a tandem mass spectrum of specific fragment ions for each peptide. Peptide sequences (and hence protein identity) were determined by matching protein databases with the acquired fragmentation pattern by the software program, Sequest (Thermo Fisher Scientific, Waltham, MA)[16]. All databases include a reversed version of all the sequences and the data was filtered to between a one and two percent peptide false discovery rate.

**diGLY remnant purification and identification by LC-MS/MS:** Synchronized parasites (3-9 hpi) were isolated as described previously[17], using 0.015% saponin, on ice, for 10 minutes. Following washes, parasite pellets were stored at -20 °C. Samples were then prepared using the PTMScan HS Ubiquitin Kit from Cell Signaling Technology (Cat. No. 59322). Following lysis, protein concentrations were measured using the BCA assay (Thermo Cat. No. 23225) and lysates were normalized to 5 mg/mL protein. The sample preparation was then completed according to the manufacturer's instructions. Peptides were then dissolved in solvent-A, separated over a C18 capillary column, ionized by electrospray, and identified using an LTQ Oribtrap Velos Elite ion-trap mass spectrometer (see coimmunoprecipitation, above).

**Immunoblotting:** Synchronized parasites were isolated on ice for 10 minutes with 0.015 % saponin. Isolated parasites were washed three times with ice-cold PBS containing a protease inhibitor cocktail (Roche cOmplete mini, Sigma Cat. No. 11836153001) before lysis in 1x Laemmli Sample Buffer (Biorad Cat. No. 1610747) containing 5% beta-mercaptoethanol, followed by heating the samples at 99°C for 10 minutes. Protein samples were separated on Mini-PROTEAN® TGX™ Precast Gels (4–20% gradient, BioRad Cat. No. 4561091) in Tris-SDS-glycine buffer. Separated proteins were transferred to a nitrocellulose membrane (Life Technologies Cat. No. IB3010-31) using an iBlot system (Life Technologies). The membrane was blocked in Intercept® (TBS) Blocking Buffer (LI-COR Biosciences Cat. No. 927-60001) for 1 hour at room temperature on an orbital shaker. Membrane-bound proteins were first probed with primary antibodies [mouse anti-*Pf*Actin 5H3 (1:100; Walter and Eliza Hall Institute of Medical Research Cat. No. 26/11-5H3-1-2)[18], mouse anti-HA (1:2000; Pierce Cat. No. 26183), mouse anti-ubiquitin mAb (FK2) (1:1000; Sigma Cat. No. ST1200), or rabbit polyclonal anti-BiP (1:5000; kindly gifted by Dr. Jeffrey Dvorin, Harvard Medical School)] in Intercept® (TBS) with 0.2% Tween 20 at 4 °C overnight. The membranes were then incubated for 45 minutes at room temperature with secondary antibodies [IRDye® 680RD goat anti-mouse (1:10,000; LI-COR Biosciences Cat

No. 926-68070) and IRDye® 800CW Goat anti-Rabbit (1:10,000; LI-COR Biosciences Cat. No. 926-32211)], washed, and imaged, then analyzed using the LI-COR Odyssey CLx Imager System (LI-COR Biosciences).

**Microscopy-based endocytosis assay (ring-stage):** RBCs were suspended in a hypotonic solution and pre-loaded with fluorescein-dextran (ThermoFisher Cat. No. D22910) as previously described[19,20]. After resealing red blood cells (RBCs) and trapping fluorescein dextran within, schizonts were added and allowed to re-invade by following the RSA setup. As the parasites (rings) grew within the fluorescein-dextran-containing RBCs, fluorescein-dextran was endocytosed and accumulated in the DV or other compartments. The parasites were then treated with 5% sorbitol to remove any remaining schizonts, closely mimicking the RSA setup. Parasites were washed once with RPMI medium and released from the RBCs using 10 pellet volumes of PBS containing 0.015% saponin. Following 3 washes with RPMI medium, parasite samples were fixed with pre-warmed 4% paraformaldehyde-0.1% glutaraldehyde in PBS for 20 minutes at 37°C. Afterwards, 2-5 µL samples were loaded onto clear slides, covered with a coverslip, and allowed to settle for up to 15 minutes at room temperature before imaging. DIC and AF488 channel images were captured using a 100X oil immersion objective on a Zeiss Axio Observer. AF488 channel images were acquired at 480 ms across all samples, with a laser intensity of 40%. Images were blinded prior to analysis. Analysis was conducted for regions of interest (ROIs) where an RBC ghost could be visualized around a green channel signal in the merged image. A global threshold was set on 8-bit images of the green channel across images taken at the same exposure settings. Area, mean gray value, and integrated density of the parasite-specific green-channel signal were measured using ImageJ (https://imagej.net/ij/).

**Coverslip IFA of late-stage parasites:** Late-stage parasites were settled on to a poly-D-lysine coated #1.5 12 mm coverslip for 30 minutes. After removing the supernatant, the parasites were fixed with 4% paraformaldehyde for 10 minutes, then washed 3 times in PBS. Cells were permeabilized with 0.1% Triton X-100 in PBS for 10 min and washed three times in PBS for 3 min. Blocking solution (3% (w/v) BSA in PBS) was added and samples were incubated overnight at 4 °C. Primary antibodies were diluted in blocking solution and added to the coverslips for 2 hours at room temperature. Coverslips were washed 3 times for 3 minutes in PBS and then incubated with secondary antibody in blocking solution for 45 minutes. Coverslips were washed three times with PBS, then mounted onto a slide with VectaShield Vibrance with DAPI. Parasites were visualized on a Zeiss LSM980 with Airyscan2 for super-resolution microscopy. A 63x objective with a numerical aperture of 1.4 was used. Dilutions for primary antibodies were 1:200 for rat anti-HA and 1:300 for rabbit anti-*Pf*Actin.

**Ultrastructure Expansion Microcopy (U-ExM):** Late segmented schizont stage *Pf*Coronin$^{WT}$-smHA or *Pf*Coronin$^{R100K/E107V}$-smHA parasites were purified by Percoll gradient centrifugation. Various age groups of parasites, as specified in the results section, were utilized for U-ExM. For experiments aimed at confirming RBC content uptake, RBCs in a hypotonic solution were pre-loaded with biotin-dextran (ThermoFisher Cat. No. D1956), as described in the endocytosis assay above. The samples were treated with pre-warmed 4% paraformaldehyde-0.1% glutaraldehyde in PBS for 20 minutes at 37 °C and prepared for U-ExM following the procedure described previously[7,21]. The following antibodies were employed for visualization: rat anti-HA for *Pf*Coronin-smHA (1:25; Sigma Cat. No. 11867423001), Sytox Deep Red for DNA (1:1000; ThermoFisher Cat. No. S11380), rabbit anti-aldolase as a marker for parasite cytoplasm (1:2000; Abcam Cat.

No. ab207494), Streptavidin-488 for parasites containing internalized biotin-dextran (1:250; Thermo Fisher Cat. No. S11223), and Alexa Fluor 405-conjugated NHS ester for protein density (1:200; ThermoFisher Cat. No. A30000). All secondary antibodies (anti-rat IgG Alexa fluor 488 Thermo Fisher Cat. No. A11006, anti-rabbit IgG Alexa fluor 555 Thermo Fisher Cat. No. A21428) were diluted 1:500 in PBS.

**Statistical analysis:** All experiments were performed with at least three biological replicates, unless described otherwise. For volcano plots of proteins from *Pf*Coronin coIPs, *p*-values were computed using unpaired t-tests. Protein quantifications (immunoblot) were compared using paired t-tests. RSA assays were compared using unpaired *t*-tests with the Welch correction. Size comparisons (e.g. parasite and invagination perimeters) were compared using unpaired t-tests. Uptake of host cell contents (endocytosis assay) was compared using unpaired non-parametric Mann-Whitney tests. The presence or absence of preDV /invagination in WT and mutant parasites was assessed using Fisher's exact test, specifically employing the Baptista-Pike method. All statistics were calculated using GraphPad Prism. No statistical methods were used to predetermine sample sizes. Researchers were blinded for analysis of the endocytic assay (uptake of host cell contents) and for all RSA assays.
